# Modular mathematical analysis of the control of flagellar Ca^2+^-spike trains produced by CatSper and Ca_V_ channels in sea urchin sperm

**DOI:** 10.1101/415687

**Authors:** D.A. Priego-Espinosa, A. Darszon, A. Guerrero, A.L. González-Cota, T. Nishigaki, G. Martinez-Mekler, J. Carneiro

## Abstract

Intracellular calcium ([Ca^2+^]_i_) is a basic, versatile and ubiquitous cellular signal controlling a wide variety of biological processes. A remarkable example is the steering of sea urchin spermatozoa towards the conspecific egg by a spatially and temporally orchestrated series of cytosolic [Ca^2+^]_i_ spikes. Although this process has been an experimental paradigm for reproduction and sperm chemotaxis studies, the composition and regulation of the signalling network underlying the cytosolic calcium fluctuations are hitherto not fully understood. Here, we used a differential equations model of the signalling network to assess which set of channels can explain the characteristic envelop and temporal organisation of the [Ca^2+^]_i_-spike trains. The signalling network comprises an initial membrane hyperpolarisation/repolarisation produced by an upstream module triggered by the egg-released chemoattractant peptide, via receptor activation, cGMP synthesis and decay. Followed by downstream modules leading to pH_i_, voltage and [Ca^2+^]_i_ fluctuations. The upstream module outputs were fitted to kinetic data on cGMP activity and early membrane potential changes measured in bulk cell populations. Two candidate modules featuring voltage-dependent Ca^2+^-channels link these outputs to the downstream dynamics and can independently explain the typical decaying envelop and the progressive spacing of the spikes. In the first module, [Ca^2+^]_i_-spike trains require the concerted action of a classical Ca_V_-like channel and a potassium channel, BK (Slo1), whereas the second module relies on pH_i_-dependent, [Ca^2+^]_i_-inactivated CatSper dynamics alone. The model predicts that these two modules interfere with each other and produce unreasonable dynamics when present at similar proportions, which suggests that one may predominate over the other *in vivo*. To assess these alternatives, several quantitative predictions were derived from each module and confronted to experimental observations. We show that the [Ca^2+^]_I_ dynamics observed experimentally after sustained alkalinisation can be reproduced by a model featuring the CatSper module but not by one including the pH-independent Ca_V_ and BK module. We conclude in favour of the module containing CatSper.

## 1 Introduction

Intracellular calcium ([Ca^2+^]_i_) plays a key role as a second messenger in a wide variety of biological processes. Its elevation above resting levels conveys diverse physiological signals and may trigger cellular processes such as programmed cell death and egg activation. Both versatility and specificity of this ion as a physiological signal lie in how it is temporally and spatially organised within a cell [1]. Calcium oscillations are widely described in both excitable and non-excitable cells, and lie at the core of several signalling processes, such as neuronal firing, embryonic cell differentiation, immune cell activation and rhythmic beating of the heart. The mechanistic basis of these oscillations have been worked out in a number of cells. A good example is the calcium induced calcium release (CICR), which is generated by an inositol trisphosphate and calcium signalling system that controls a host of biological processes [2], for which its operation has been understood with the help of mathematical models [3, 4]. Furthermore, in olfactory neurons, calcium influx is regulated directly by cyclic nucleotides [5, 6].

The guidance of sea urchin spermatozoa towards their conspecific eggs during broadcast spawning events is orchestrated by [Ca^2+^]_i_-spike trains elicited by the chemoattractant sperm activating peptides (SAP) released by the eggs. These spike trains are decoded into a series of acute turns followed by straight swimming episodes that define the trajectory of the sperm cell. Preventing calcium influx from the external medium will abrogate the cell capacity to orientate itself in space. The sequence of spikes can be triggered by SAP in sperm that are fixed, showing a typical temporal organisation, characterised by a gradual decrease of the amplitude of the spikes and progressive increase of the interspike intervals (Fig. 3).

The signalling network that mediates calcium-steered chemotaxis in sea urchin sperm has been an object of intense study, and several decades of experimental research have implicated diverse channels and molecular components [7]. It is now clear that the first step leading to Ca^2+^ influx is an increase of intracellular cGMP concentration, which is synthesised in response to the natural ligand. Experiments in which the receptor activation was bypassed by releasing a caged analogue demonstrated that cGMP is sufficient to trigger the signalling network downstream. The rise in cGMP leads to membrane hyperpolarisation by opening a cGMP-gated K^+^-channel (KCNG). How the hyperpolarisation is then transduced into Ca^2+^ flux dynamics is not fully established. It has been proposed that the membrane potential shift to more negative values enables a depolarising cationic current by sperm specific hyperpolarisation-activated cyclic nucleotide-gated channels (spHCN), and removes the inactivation of voltage-gated calcium channels (Ca_V_), which will then further depolarise the membrane. The depolarisation of the membrane turns off spHCN and Ca_V_, allowing hyperpolarisation fluxes to predominate, repeating the cycle. This cycle would be the basis of the [Ca^2+^]_i_-spike train, the temporal organisation of which could be modulated by Ca^2+^-activated Cl^-^ and K^+^ channels. Another possibility is that the cGMP-driven hyperpolarisation, by activating a Na^+^/H^+^ exchanger that extrudes protons, rises the pH_i_ which in turn elicits Ca^2+^ influx by a pH-dependent Ca^2+^ channel, such as CatSper. This is a sperm specific calcium channel with unique features that separate it from the classical Ca_V_ family, including modulation of voltage-gating by pH_i_ and inactivation by Ca^2+^. These features open the possibility for another negative feedback underlying [Ca^2+^]_i_ oscillations. The pH_i_ rise elicits the opening of CatSper that drives an upsurge in [Ca^2+^]_i_ and membrane depolarisation, which in consequence inactivate respectively CatSper and the exchanger. The cycle reinitiates once the membrane becomes again hyperpolarised.

The evidence for the two types of calcium channels proposed to participate in SAP-activated [Ca^2+^]_i_-spike trains is summarised in Table 1. Most evidence is indirect and based on pharmacological approaches, which attempt to implicate channels using blocking and activating drugs, that often are not specific and therefore have off-target effects. Genetic engineering techniques that would allow definitive conclusions on the necessity and sufficiency of the posited channels (e.g. null mutants) are not available. Therefore, it remains elusive whether a single or multiple types of calcium channels play an active role in sea urchin sperm signalling.

**Table 1.** Summary of experimental evidence on the main calcium channels proposed in the pathway activated by SAPs. ^a^It is worthwhile to mention that out of the pharmacological blockers listed, none of them are 100% specific to the associated channels. ^b^Antibody raised against a polypeptide of the *α* subunit in rat, which also recognises the homologous version in mouse and human sperm. ^c^Antibody raised against pore-forming subunit polypeptides of *A. punctulata* [14] and *S. purpuratus* [15]. ^d^ It is of note that direct electrophysiological recordings for this ion channel have not been achieved in mature sea urchin spermatozoa compared with the homologous complex in mouse, human and macaque spermatozoa [16].

| $\text{Ca}^{2+}$ channel | Experimental evidence on sea urchin sperm | |
| --- | --- | --- |
|  | Molecular biology | Pharmacology<sup>a</sup> |
| LVA (T type) | mRNA in testis tissue[8] | Nickel [9]<br>Nimodipine [10, 11] |
| HVA (L type) | mRNA in testis tissue,<br>immunolocalisation<sup>b</sup>[12] | Nimodipine [12]<br>Nifedipine [12]<br>Verapamil [13] |
| CatSper<sup>d</sup> | mRNA in testis tissue,<br>immunolocalisation<sup>c</sup>,<br>mass spectrometry[14, 15] | Mibefradil [14]<br>MDL12330A [14]<br>NNC 55-0396 [15] |

Mathematical modelling may help to explore alternative hypotheses that cannot be directly addressed and disentangled experimentally. The signalling network elicited by Speract in sea urchin sperm has been modelled using the discrete formalism by [13, 15, 17]. While the initial discrete network models featured only Ca_V_s, a subsequent attempt to gain insights into the effect of multi-target niflumic acid suggested that pH-dependent CatSper would play a key role in Ca^2+^ oscillations, and that the calcium-dependent potassium channel (BK) would further modulate the spike trains [17]. The inherent limitations of discrete models in describing concentrations and kinetic details hinder the conclusions that can be drawn from comparison with quantitative experiments. A continuous model was used to identify the core components of the signalling network [18], implicating Ca_V_ channels, but the model was not directly compared to experimental data.

In the present work, we explore which putative calcium channels may mediate the Ca^2+^ response to SAP by describing the network using coupled ordinary differential equations that can fully harness the quantitative and kinetic information available in experimental data. We asked which channels may intervene, and how they may control the relevant dynamical features of the observed [Ca^2+^]_i_-spike trains. Our aim is to contrast the different hypotheses by their quantitative implications in order to identify critical properties that can be used to experimentally disentangle them.

## 2 Materials and Methods

### 2.1 The general modelling framework

Our strategy is to describe the dynamics of the SAP-activated signalling pathway in order to understand how the [Ca^2+^]_i_ fluctuations are controlled by different molecular components. A system of ordinary differential equations (ODE) describes the dynamics of the membrane potential and of the intraflagellar concentrations of cGMP, protons and Ca^2+^. These four essential variables depend on the activities of different flagellar channels and enzymes, the dynamics of which are also described by ODE (Sec. A1). The four variables are the main observables in the model and offer the means to assess its predictions by comparison with experimental data.

#### 2.1.1 Membrane potential

The dynamics of the membrane potential, *V*, is described with a Hodgkin-Huxley formalism [19]. The temporal derivative of *V* is defined as a sum of ion current densities normalised by the flagellum specific capacitance, according to Kirchoff’s law of charge conservation:

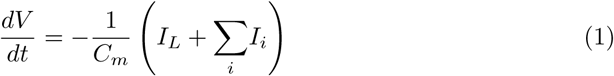

Each current, indexed by *i*, is associated to a given channel type to be chosen from a set of ion channels, i.e. *i* ∈ {KCNG, spHCN, Ca_V_, BK, CatSper}. The current terms *I_i_* are defined according to Ohm’s law:

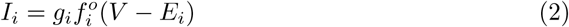

where the *E_i_* is the reversal potential of the channel *i*, 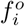 is the fraction of open channels conducting current and *g_i_* is an effective conductance of the channel defined as the product of the channel unitary conductance by the channel density in the flagellum membrane.

Following the Hodgkin-Huxley tradition, we define a leakage current term *I_L_* = *g_L_*(*V − E_L_*) that allows to ensure that the resting membrane potential *V*̂ = *E_m_* is asymptotically reached by imposing the following constraint on *E_L_*, given the parameters and variables *f_i_* at equilibrium:

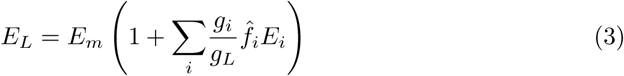

The leakage current is interpreted as the net current arising from all the channels and transporters that are not explicitly described in the model.

As a remark on notation, in this article a hat-decorated variable *x*^ refers to the equilibrium value of the variable *x*.

#### 2.1.2 Cyclic nucleotide concentration

The concentrations of the nucleotides cGMP and cAMP increase in the sperm flagellum in response to SAP stimuli. These nucleotides affect the permeability of several nucleotide-gated ion channels, in turn changing membrane potential. Considering that cGMP is the nucleotide showing the highest changes after SAP binding and that the experimental release of its analogues in the flagellum is sufficient to produce a dose response that closely mimics the response elicited by the physiologic ligand, we restricted our analysis to this nucleotide, neglecting the eventual role of cAMP.

The intraflagellar concentration of cGMP is denoted by *G* and its dynamics described by the following differential equation:

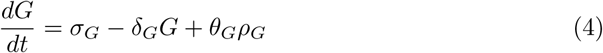

In this equation, it is assumed that the activities of general guanylate cyclases and phosphodiasterases lead to a constitutive turnover of cGMP, with a constant source *σ_G_*, and turnover rate *δ_G_*. In the absence of SAP signals, this basal turnover results in a cGMP at rest stationary concentration *G_r_* = *σ_G_/δ_G_*. The basal turnover is assumed to be perturbed by the additional production of cGMP due to active SAP receptors, described by the term *ρ_G_* (Sec. A1.1).

*G* was compared with measurements of total cGMP by radioimmunoassay in bulk cell experiments previously reported [20]. Since *G* represents the effective intraflagellar concentration of cGMP available to bind to KCNG channels, it was necessary to introduce a scaling factor for the term *ρ_G_*. This factor, *θ_G_*, consists of the product of flagellar volume (*υ_f_*), cGMP buffering capacity in the flagellum (*B_G_*) and Avogadro constant (*N_A_*). In this way, only a fraction of the total cGMP synthesised by the active receptors participates in the signaling response and the *G* signal is kept within the nm sensitivity range reported for KCNG channels [21] (Sec. A1.2).

#### 2.1.3 Proton concentration

The permeability of some channels is affected by cytoplasmic concentration of protons. The intraflagellar proton concentration, denoted by *H*, is described by the following differential equation:

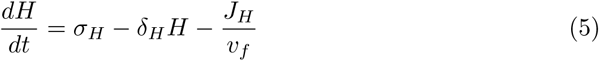

which assumes that there is a source of intraflagellar protons, *σ_H_*, a basal turnover with rate *δ_H_* and an efflux *J_H_* mediated by the activity of the sodium/proton exchanger (Sec. A1.4). The parameter values were constrained by imposing the basal proton concentration to be 7.9 × 10^*−*8^ m (pH_i_=7.1, Tab. A1), obtained by solving *H* for equilibrium. The relative changes of the variable *H* in time were further compared with pH-sensitive, fluorescent probe signals measured in bulk populations [22] or in individual cells [23].

#### 2.1.4 Intraflagellar Ca^2+^ concentration

The intraflagellar Ca^2+^ concentration, represented by the variable *C*, is the main observable of our study. Its dynamics is described by the following differential equation:

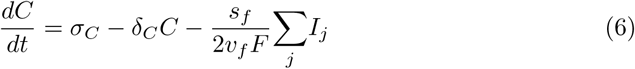

with *j* ∈ {Ca_V_, CatSper}. Just as in the case of the previous components, we assume that different sources and sinks of intraflagellar Ca^2+^ result in a net constant production rate *σ_C_* and a basal turnover rate *δ_C_*. The [Ca^2+^]_i_ dynamics is further controlled by the activities of specific Ca^2+^ channels, described by the third term in the equation as the sum of ionic current densities scaled by the ratio of flagellum membrane area over the product of ion charge of a Ca^2+^ ion, flagellar volume and Faraday constant. The list of currents to be considered will depend on the scenario being analysed.

The variable *C* was compared to measurements by Ca^2+^-sensitive fluorescent probes, either in individual cells or in populations [10, 20, 24]. Individual cell measurements were used to analyse the temporal structure and the magnitude of the Ca^2+^ spikes, while population measurements gave only information about the envelop of the spike train.

### 2.2 Computational implementation and parameter estimations

The systems of differential equations were solved numerically with Mathematica 11 (Wolfram Research, Inc., Mathematica, Version 11.0, Champaign, IL), XPPAUT 8 [25] or R (using the package `deSolve`[26]). XPPAUT 8 was used for the bifurcation analysis (see below). Mathematica was further used to fit the model to experimental data using the `Nonlinearmodelfit` function with the optimisation method `Differential Evolution` with constraints. In the analysis, we defined several model variants based on subsets of the variables and equations presented above and in section A1. Some of these defined modules were used to estimate the parameters and some were used to analyse specific properties.

The upstream module fitting was done in two parts:

- The state space comprised by the variables {*S*, *R_H_*, *R_L_*, *G*}, which includes the reactions involving the SAP-receptor (eq. 12) and the cGMP dynamics (eq. 4), was fitted to kinetic data of cGMP measured in bulk cell populations [20]. Data points were read from the indicated publication and are expressed in units of pmol per 10^8^ cells. The cGMP kinetics obtained with the different SAP concentrations used in [20] were fitted simultaneously with the model (Fig. S1). For this fitting, the parameter *θ_G_* (described in the section 2.1.2) was not included.
- *•* The remaining variables of the module 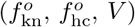, which involve the early electrophysiological changes caused by SAP (i.e. hyperpolarisation and repolarisation due to the opening of KCNG and spHCN, respectively), were used to fit the gating parameters of the ion channels. The spHCN parameters were either extracted or calculated from [27–30], while those of KCNG were based on the values reported in [8, 21, 31]. Fixing these values, the conductance density parameters of these channels were calibrated together with *B_G_* (included as a free parameter in the scaling factor *θ_G_*), by fitting the dynamics of *V* with the kinetic data from [8], which comprise the early response of the membrane potential change. Because this module is only intended to explain the early transient behaviuor of *V* in terms of these two channels, we only considered the data of the first second of the time series.

In order to calibrate this module, we assumed that the overall dynamics under both single-cell and population regimes can be comparable by using a scaling factor; this depends on the flagellar volume term to convert the cGMP signal into the effective concentration capable of exerting a physiological effect at the single cell level (see parameter *θ_G_*, Sec. 2.1.2).

To study the dynamics of intraflagellar calcium, we set out to explore the possible contribution of different channels under two main scenarios.

- In one scenario, the model features the Ca_V_ + BK module and has the state space 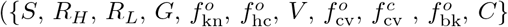. The values of the Ca_V_ channel parameters were adjusted based on the characteristics of low-voltage-activated T-type Ca^2+^ channels [32], for which there is evidence suggesting their presence in sperm [8, 10] (Tab. 1).
- In the alternative scenario, the model features the CatSper module and has the following state space 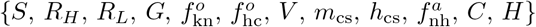. The CatSper gating parameters were set such that its Ca^2+^_i_ and pH_i_ sensitivity range is placed within the physiologically expected values in reported natural SAP responses (Fig. S3).

### 2.3 Mixed scenarios parametrisation

We also analysed mixtures of Ca_V_+BK and CatSper modules, in the most extended model with state space 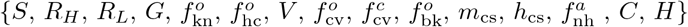. We introduced a parameter *θ* to explicitly weigh the contribution of each combination of channels. The latter parameter modulates CatSper, Ca_V_ and BK in the following way:

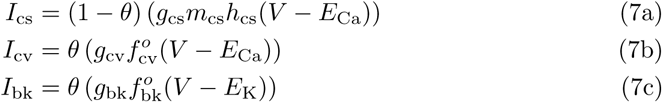

From the above equations, we can express a ratio of effective maximal conductance of Catsper with respect to that of Ca_V_:

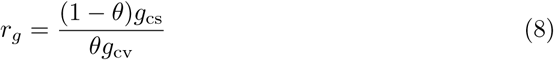

Always taking the CatSper module as a starting point, the analysis of mixed scenarios was done by adding Ca_V_ and BK channels in three ways: a) Ca_V_ only, b) BK only, and c) Ca_V_ + BK.

Because the parameters that control the equilibrium state at rest differ between the two alternative modules (*g_L_* and *δ_C_*, see Tab. A1), to ensure a consistent recovery of the behaviours predicted by each module in extreme cases (*θ* = 0 and *θ* = 1), these parameters were rescaled in a *θ*-dependent manner according to the following formulas:

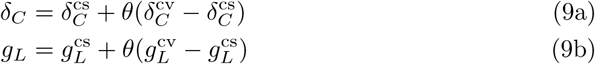

### 2.4 Resting state and initial conditions

Initial conditions of the gating variables of KCNG, spHCN, Ca_V_, BK and sNHE, as well as parameters *σ_C_*, *E_L_* and *δ_H_*, were all calculated by solving the system at equilibrium in the absence of SAP, that is, setting each differential equation to 0 under the condition *S* = 0. Physiological observables *G*, *C*, *H* and *V* were initialised to their respective resting values, as reported in the literature (Tab. A1).

The parameters governing sNHE activation were chosen such that approximately 40% of exchangers were active at the resting state, just as inferred from experimental data in activated sperm without any chemoattractant stimulation [33, 34], whereas the maximum effective proton flux by sNHE, *J*_max_, was fitted to limit the pH_i_ up to ~ 7.7, after physiological stimulation by SAP [35, 36].

## 3 Results

### 3.1 A modular signalling network

The potential signalling network under study here is illustrated in Fig. 1. The signalling network is organised in three modules: an upstream module, which includes the ligation of the receptor by SAP and the early transient drop of membrane potential *V* generated by the cGMP response, and two alternative downstream modules with the potential to generate the Ca^2+^ oscillatory response. The Ca_V_+BK module includes voltage-dependent Ca^2+^-channels and Ca^2+^-dependent K^+^-channel BK; while the CatSper module is composed of the voltage-, pH_i_- and Ca^2+^-dependent channel CatSper, the intraflagellar proton concentration and the voltage-dependent sodium-proton exchanger.

**Figure 1.**
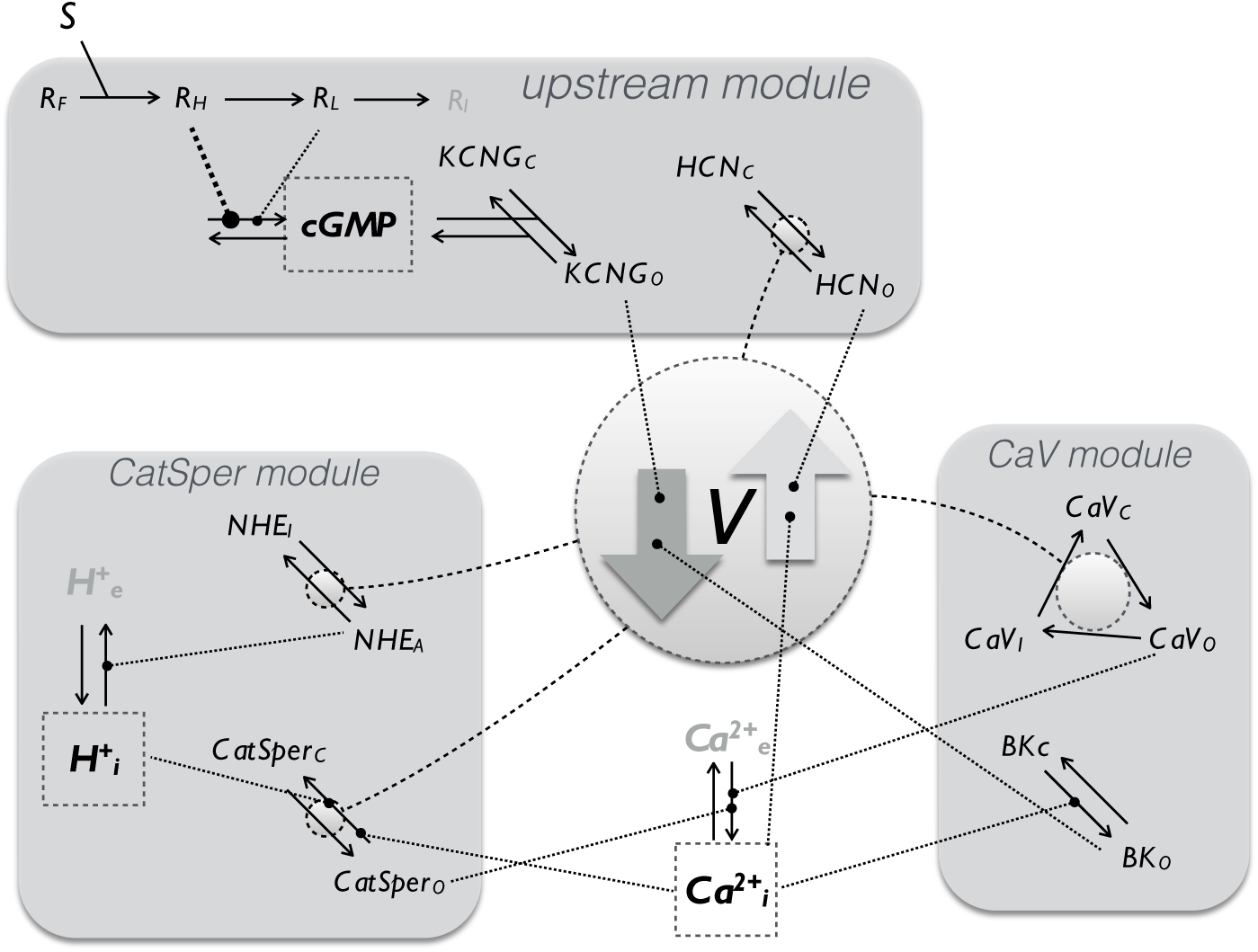
The modular organisation of the signalling networks transducing SAP signals to Ca^2+^-spike trains. The structure is separated into 3 modules linked by the membrane potential variable (*V*). In the upstream module (described by the variables 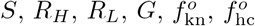 receptors (*R*) bind SAP molecules (*S*) and transit irreversibly through three states, each with less guanylate cyclase activity: High (*R_H_*), low (*R_L_*) and inactive (*R_i_*); the cGMP synthesised by these receptors opens KCNG channels, (KCNG_O_), which conduct a potassium current that hyperpolarises *V* (arrow downwards). The hyperpolarised levels of *V* promote the opening of spHCN channels (HCN_O_), which exert the opposite action on *V* when conducting a cationic inward current. Two alternative modules are presented to explain the calcium oscillation trains: one that includes classic voltage-dependent Ca^2+^ channels and BK channels, and another that considers CatSper, sNHE exchangers and proton concentration. Note in the module Ca_V_, that the calcium channel is purely voltage dependent, whereas CatSper has threefold regulation, namely Ca^2+^, H^+^ and *V*.

The modular structure facilitated the analysis of the model and addressing the question on which sets of channels control the Ca^2+^-spike trains elicited by SAP stimulus. Considering that all the components of the upstream module are independent of the variables downstream, we studied it isolated from the remaining dynamics. In so doing, we ensured that this upstream module accounted quantitatively for experimental time series of membrane potential and cGMP in response to SAP (Sec. 3.2). Once the values of the parameters of this module were obtained, they remained fixed in the subsequent analyses of the dynamics of the downstream components. In these analyses, the modules featuring Ca^2+^-channels were constrained such that given the input of the upstream module, they generated an output that was semi-quantitatively identical to the observed Ca^2+^-spike trains, i.e. these two putative modules were constrained by two independent data sets upstream and downstream. Hence, by coupling the downstream Ca_V_+BK and CatSper modules to this upstream module, we show in Sec. 3.3 that either module can independently account for the observed Ca^2+^-spike trains patterns.

### 3.2 Upstream module of the SAP-activated response: cGMP and hyperpolarisation signals

cGMP is the core component of the upstream module of SAP signal transduction. It raises in response to SAP-induced guanylate cyclase activity and activates KCNG channels that in turn lead to hyperpolarising currents. *A. punctulata* sperm stimulated with the SAP Resact exhibit a rapid cGMP response that peaks before ~400 ms and rapidly drops to a pseudo plateau that slowly decays over several seconds, as observed by stopped-flow kinetics [20]. This biphasic dynamics imposed the assumption that SAP receptors transit irreversibly through three forms with different levels of associated guanylate cyclase activity (Fig. 1).

Sufficient data is available in the literature to calibrate this module. We first fitted receptor activation and the cGMP signal to cGMP measurements in bulk cell assays, wherein sperm were stimulated with pulses of uncaged SAP along a concentration range spanning several orders of magnitude [20]. This fitting (Fig. S1) led to the values of the kinetic rate constants for receptor state transition, cGMP synthesis and decay (kinetics illustrated in Fig. 2 for a SAP concentration that saturates the *V* response [8]). Before calibrating the voltage and ion channel dynamics, and in order to afterwards calibrate the spiking module, we used appropriate scaling factors to extend model solutions towards single-cell regimes by converting total cGMP quantities into an effective concentration capable of opening KCNG channels in a sperm flagellum. By adjusting the density of KCNG and spHCN channels to membrane potential from sperm population data [8], we found values that reproduce the early transient drop in membrane potential and the early recovery (Fig. 2). To illustrate the separate contribution of each channel, we showed a reference case where KCNG is the only channel present (dashed line in Fig. 2) and, as expected, its opening alone drives membrane potential towards *E*_K_ ≈–80 mV (potassium Nernst potential). The latter reference value sets a lower limit to the *V* response that is well below the value reached at the minimum *V* produced by the KCNG+spHCN case. It is worth noticing that under this scenario, in which only these two channels are assumed to be present, it is not possible to find a given combination of channel densities such that the spHCN current alone counterbalances the hyperpolarising current carried by KCNG, so as to bring membrane potential back to resting levels without abrogating the hyperpolarisation pulse. Therefore, full recovery of voltage requires the additional activity of other type of ion channels with depolarising currents after these initial events (e.g. calcium channels).

**Figure 2.**
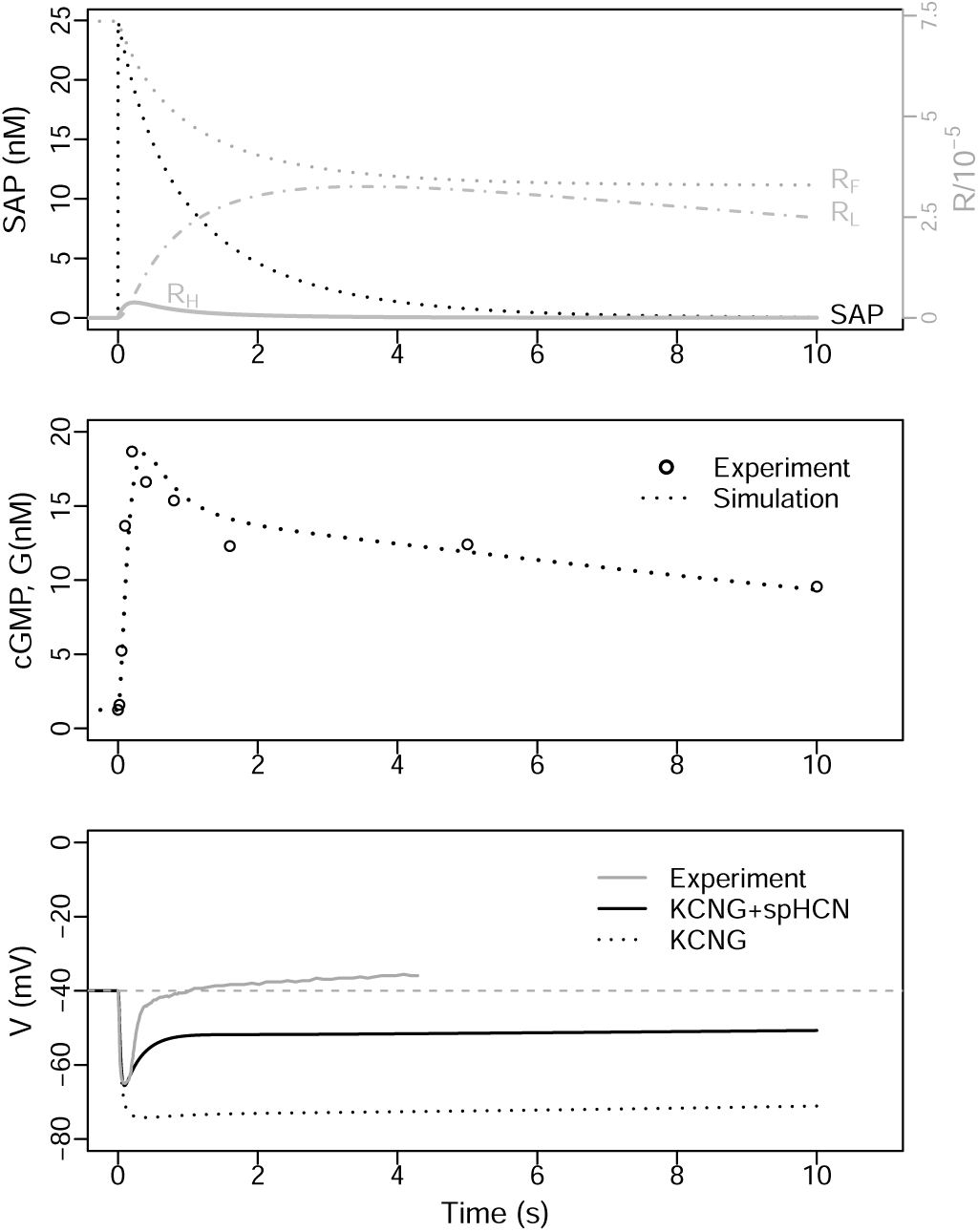
Numerical solution of the upstream module (state space 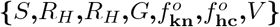) overlaid with experimental data obtained in bulk. Fitting was done with data from cell populations stimulated at *t* = 0 with a SAP pulse *S*_0_ =25 nm (data points read from graphs in publications [20] and [8] for cGMP and and *V*, respectively). In the top graph, receptor activation dynamics and SAP consumption are shown. The middle graph displays *G* dynamics with units of total amounts. In the last graph, we show two separate cases: KCNG+spHCN (solid black) and KCNG only (dotted).

### 3.3 Two candidate signalling modules can explain semiquantitatively the structure of the Ca^2+^-spike trains

Two alternative signalling modules, labelled Ca_V_+BK and CatSper, can explain the characteristic envelop and increasing delay of the Ca^2+^-spikes when coupled alone to the upstream module (see Fig. 1). These modules are composed of specific combinations of ion channels and transporters. These combinations were selected by their capacity to produce a series of Ca^2+^-spikes that recapitulates semiquantitatively the ones observed experimentally, given the input cGMP kinetics used to calibrate the upstream module (Fig. 2). In other words, the numerical solutions of these modules lead to [Ca^2+^]_i_-spike amplitudes and intervals between consecutive spikes that lie within the relative range of the spikes observed in individual sperm cells of *S. purpuratus* stimulated with the SAP Speract (Fig. 3). The two modules and their properties are described separately in the next two sections.

**Figure 3.**
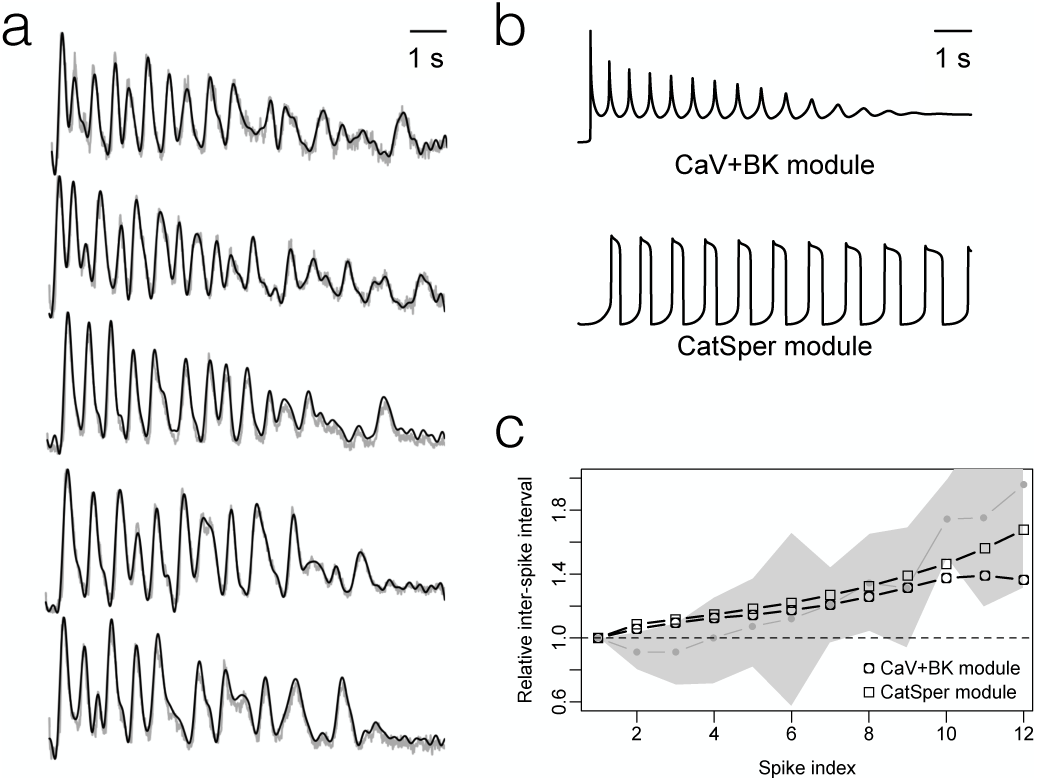
[Ca^2+^]_i_-spike trains produced by the sperm activating peptides in sea urchin sperm flagellum and its modelling. A) Examples of [Ca^2+^]_i_ time series measured with fluorescent probes in flagella of *S. purpuratus* individual sperm bound to a coverslip following uncaging of Speract analague by UV light. B) Numerical solutions of the model featuring either the Ca_V_+BK module or the CatSper module, coupled to the upstream module. C) Temporal organisation of the interspike intervals observed experimentally (the median ant interquartile range are represented by the gray dots and the gray band, respectively) and in the numerical solution of the Ca_V_+BK and CatSper modules (black symbols and lines)

#### Ca_V_+BK module and its dynamical properties

The first Ca^2+^ channel to be implicated in SAP signalling in sea urchin sperm literature was the Ca_V_ channel and following this chronology we will start the analysis of its module. The structure of the module is depicted in Fig. 1 and features the Ca_V_ channels and the Ca^2+^-dependent BK potassium channels. In our model, Ca_V_ channels transit irreversibly through three forms – inactive, closed and open – defining an ordered cycle, in which the transition rates are controlled by voltage. Only in the open form Ca_V_s allow inward Ca^2+^ currents that tend to depolarise the membrane. Provided that the transition rates between these forms are of similar order of magnitude, these channels alone give rise to a limit cycle, which as we will see has a constant period and relatively low amplitude (Fig. 5B) as compared to those of Fig. 3. The fact that the period predicted by these channels alone is constant would be incompatible with the progressive increase of the interspike intervals observed experimentally. This disagreement with the observations called into action the other channel in the module, BK, which once coupled with the Ca_V_ introduces a progressively increasing delay in the interval between the peaks and also increases the amplitude of the Ca^2+^ spikes.

The numerical solution for the variables in the submodel including the Ca_V_+BK module 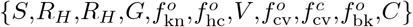 is shown in Fig. 4 for the reference parameters. The uprise and slow decay of cGMP produced by the upstream module leads to Ca^2+^-spike trains with an envelop that follows closely the cGMP values (top) and increasing interspike intervals. The initial transient hyperpolarisation (bottom) driven by the opening and currents of KCNG (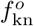 and *I*_kn_) precedes and triggers the first peak of [Ca^2+^]_i_ (*C*). The nadirs of the [Ca^2+^]_i_ oscillations are well above the basal concentration of this cation and this minimal plateau of about 500 nm is maintained until the Ca^2+^ activity vanishes at about 17 s. A similar oscillatory behaviour with similar period but different phases is observed on the fractions of open channels (second row) and the respective currents (third row), as well as the membrane potential (bottom).

**Figure 4.**
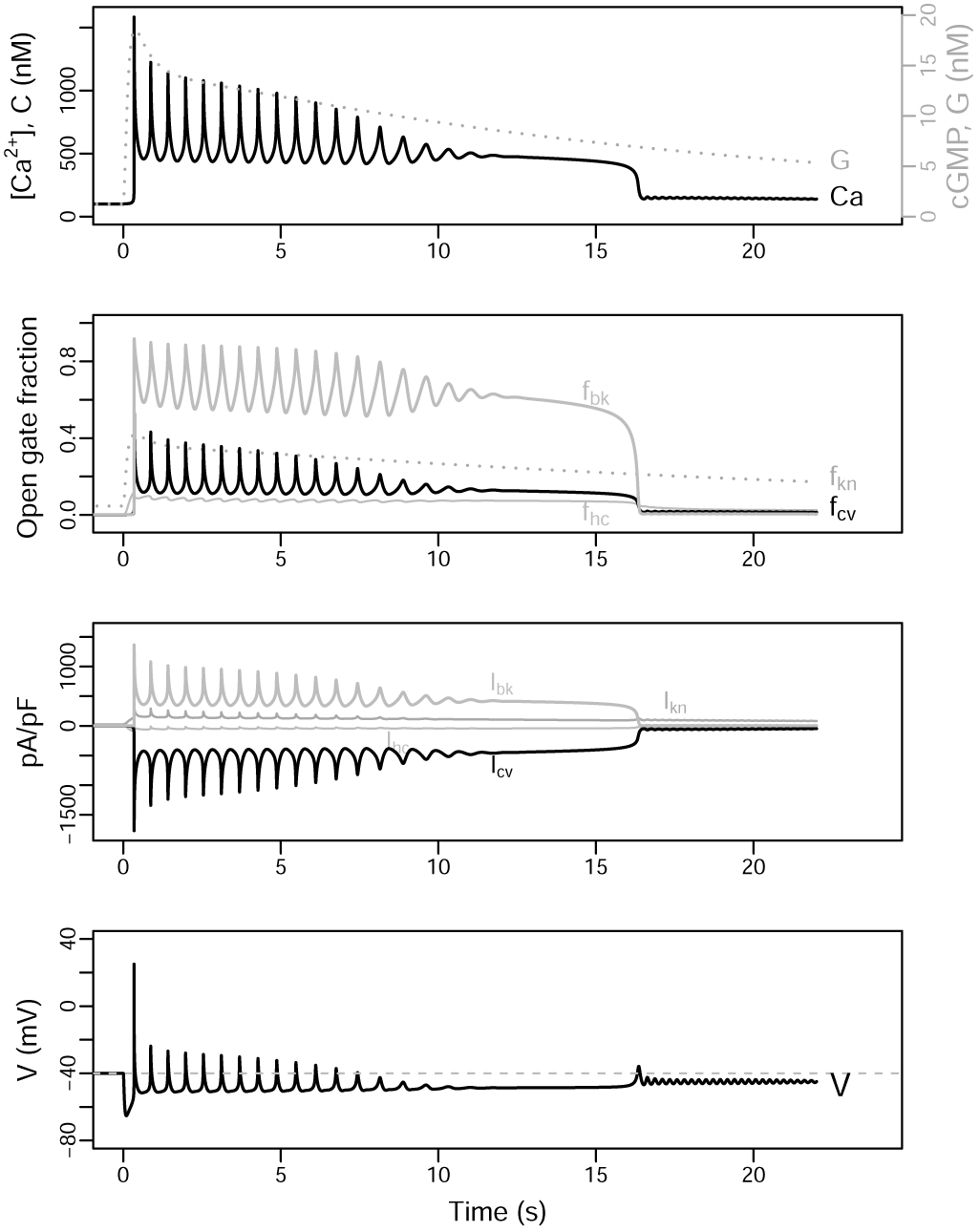
Dynamics of the model featuring the Ca_V_+BK module (state space 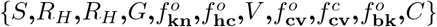). As in Fig. 2, an initial stimulus *S*(0) = 25 nm is used. From top to bottom, the numerical solutions obtained with the parameters listed in Tab. A1 are shown for: *C* together with *G* (rescaled so that their respective basal values coincide), open channels fraction, ionic currents and membrane potential.

The temporal dynamics of the [Ca^2+^]_i_ fluctuations predicted by this module can be easily understood qualitatively. The hyperpolarisation induced by KCNG enables the transition from inactive to closed forms of the Ca_V_ and the spHCN-mediated increase in membrane potential opens these closed channels. The open Ca_V_ channels will overcome the hyperpolarising current driven by KCNG and thus depolarise the membrane as Ca^2+^ ions flow in. The depolarisation of the membrane leads to the inactivation of the Ca_V_ channels with the hyperpolarisation currents predominating again. At the hyperpolarised membrane potentials the inactive channels will again transit to closed form, which eventually will open as described above, reinitiating the cycle. Intraflagellar calcium, by activating the BK channels, adds a strong hyperpolarising current to that of KCNG. This couples the net hyperpolarising currents to the amplitude of the previous Ca^2+^-spike, in such a way the higher the [Ca^2+^]_i_ peak, the higher the BK-driven hyperpolarising current will be.

The envelop of the [Ca^2+^]_i_ spikes follows closely that of the cGMP levels upstream. This led us to speculate that the time course of cGMP might offer a way to understand the fluctuations envelop. To gain a better quantitative insight, a variant of the model was made wherein *G* instead of a variable became a constant parameter, thus ignoring the upstream variables and reducing the state space (denoted 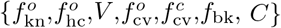). In this simplified model, the qualitative changes of the levels of [Ca^2 +^]_i_ and their periods present in stationary states were analysed as a function of the *G* parameter. This is known as bifurcation analysis in dynamical systems theory, which is useful for characterising the global properties of a system of differential equations, in particular the dependence of asymptotic behaviours on a given parameter of interest (bifurcation parameter). These behaviours can correspond to attractors of fixed point (invariant in time) or limit cycles (stationary periodic), which in turn can be stable or unstable, among others. Wherever there is a qualitative alteration of these behaviours by changing the parameter of interest, it is said that a bifurcation has occurred.

When *G* is varied in this subsystem, the branch diagram shows 5 bifurcation points (Fig. 5A, labeled from I to V). In I, by means of a Hopf-type bifurcation, as *G* increases above the basal level, the system moves from a [Ca^2+^]_i_ stable stationary state to a limit cycle. When it reaches II, the limit cycle disappears giving rise again to a stable stationary equilibrium by means of an inverse Hopf-type bifurcation. Further on in III, the system undergoes an inverse saddle-node bifurcation, defined over *G* increments, in which a stable equilibrium state coalesces with an unstable one. In this way, in traversing the branch over the stable equilibrium state, the system goes to an unstable state. Now, when advancing on this last unstable equilibrium, this turns to a stable one by means of the direct saddle-node bifurcation (IV). The presence of points III and IV suggest the possible existence of a cusp bifurcation defined in a space of higher dimension, which is corroborated in panel B of the same figure. Finally, the last stable equilibrium state, again by means of a direct Hopf-type bifurcation, gives rise to another limit cycle.

**Figure 5.**
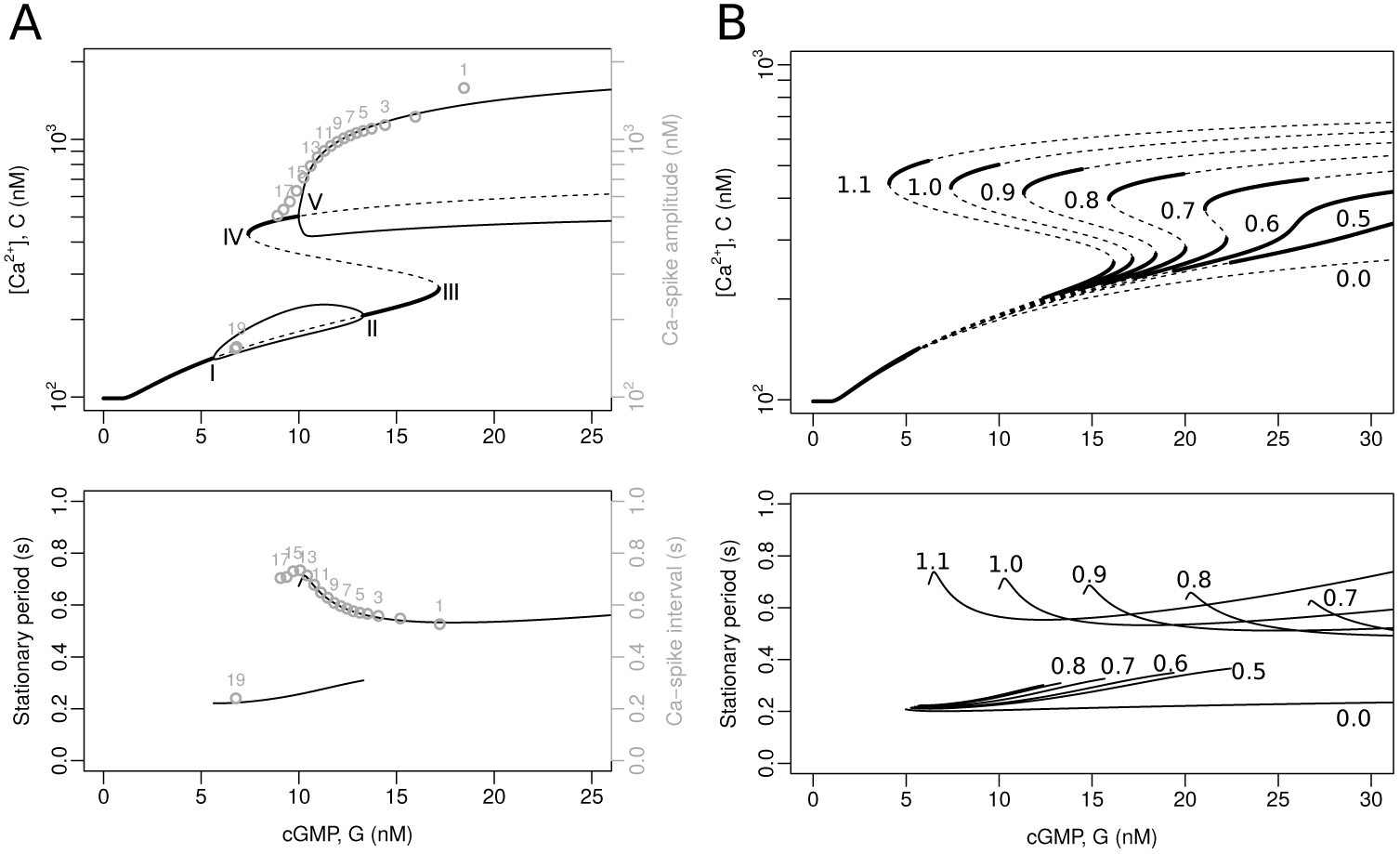
Bifurcation analysis of the model featuring Ca_V_+BK. In this analysis, cGMP is a constant input, i.e. *G* is the bifurcation parameter and the upstream variables are ignored, thus reducing the state space to 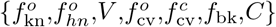 Bifurcation diagrams calculated with XPPAUT [25] using the reference parameters (Tab. A1). Top: the graphs show the *C* variable as a function of the input parameter *G*. The thick continuous lines are stable equilibria, the dashed ones indicate unstable equilibria, the thin continuous lines are the maxima and minima of stable limit cycles. The bifurcation points are marked with Roman numerals (I to V). The numbered gray circles indicate the value of the peaks obtained by numerically solving the complete model (see Fig. 4). Bottom: the solid lines represent the period of stable limit cycles as a function of the constant input *G*. The numbered gray circles correspond to the interspike interval in the numerical solutions, as in the top graph. B) Bifurcation diagrams parameterised by BK conductance density (*g*_bk_). Top and bottom: the lines are as in the graphs in (A); for clarity, the minima and maxima of the limit cycles are omitted. The numbers in the range of 0.0 to 1.1 are the fold change factors multiplying the reference value of *g*_bk_ (e.g. the curves labelled 1.0 coincide with those of (A), whereas the curves marked 0.0 correspond to a cell without BK channels)

The period of the limit cycle in the upper branch increases as the concentration of cGMP decreases (Fig. 5B, bottom), but it is not the case for the limit cycle in the lower branch, in which the period shows an opposite trend, with a less pronounced degree of change. In addition, there is a considerable difference between the two limit cycles in terms of their amplitude, demarcated by the minima and maxima of calcium levels in the bifurcation diagram.

To better understand the modulating role of the BK channel on calcium dynamics in a Ca_V_-dependent scenario, bifurcation diagrams were calculated within the *G* range considered in Fig. 5A, for different density values of BK (Fig. 5B). In B, unlike the upper panel of the figure 5A, the limit cycles depicted by the lower and upper lines are not shown for clarity; their respective unstable steady-state are shown instead (dashed lines). In the bifurcation analysis, the extreme case where *g*_bk_ = 0, that is, the variant of the model that only presents Ca_V_, predicts the existence of a single branch for the calcium concentration with a limit cycle whose envelop is controlled by cGMP levels but has a constant period of approximately 0.2 s (see the lines labeled 0.0, in both panels of Fig. 5). The gradual addition of BK reveals how bistability is generated (Fig. 5B), and exhibits the appearance of an upper limit cycle within the range of *G* values observed in Fig. 5A. The behaviour of the periods associated with the different values of *g*_bk_ is shown in the lower panel of the Fig. 5B. In a three-dimensional representation, with *g*_bk_ as one of the axes, the bifurcation pattern is a cusp with mounted limit cycles. The presence of the new upper periodic attractor allows the nadir of the [Ca^2+^]_i_ fluctuations to be placed on a plateau above the basal level, and both their amplitude and increasing interspike interval to approach the physiological response. This implies that the aforementioned properties arise from the coupling of the Ca_V_ and BK channels.

The parameters have been chosen such that as the cGMP values transiently exceed the bifurcation point, the system is forced to oscillate in the upper limit cycle and fall into the lower limit cycle after the decrease in cGMP causes the system to pass through the saddle-node bifurcation (IV). This explains the abrupt [Ca^2+^]_i_ drop and the low amplitude oscillations that persist afterwards, which are particularly visible in the numerical solution of *V* (Fig. 4, bottom). If the value of *g*_bk_ were increased above the reference value used in panel A (e.g. by multiplying it by a factor of 1.1), the saddle-node bifurcation would occur at a lower *G* value and these low amplitude fluctuations would not be revealed (see Fig. 5B).

Considering the bifurcations of the model obtained with cGMP as control constant, the behaviour of the full model is easy to understand: as SAP binds to its receptor, a sustained yet decreasing cGMP activity leads the system to approach asymptotically the attractors of the model with constant cGMP, such that the system displays fluctuations whose spike amplitude and interspike interval tend to the amplitude and the period of the attractors (compare the numbered gray circles corresponding to the solution of the full system, which takes into account the temporal variation of *G* as in Fig. 4, to the asymptotic limit cycles for different fixed cGMP values in Fig. 5A).

It is important to note that at intermediate levels of cGMP, there is a range of values where limit cycles coexist. From what has been reported in the literature, only the dynamics compatible with the upper limit cycle has been experimentally observed. Presumably, even if the small oscillations of the lower limit cycle were present, they would be masked by measurement errors.

#### CatSper module and its dynamical properties

We now consider the alternative module whose central component is CatSper, the Ca^2+^ channel that has recently been implicated in SAP signalling in sea urchin sperm. We assumed here, that CatSper opening relies on two independent gates: one modulated by both pH_i_ and voltage, and the other one responsive to intracellular calcium. The fraction of open channels is given by the product of the fractions of these two open gates (eq. 34). The module includes also as variables the intraflagellar concentration of protons, *H*, and the fraction of active forms of the voltage-sensitive Na^+^/H^+^ exchangers, 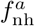. In this way, the model that results from coupling this module with the upstream module has the state space 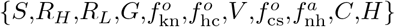. The numerical solutions obtained under the same conditions as in the alternative model are shown in Fig. 6.

**Figure 6.**
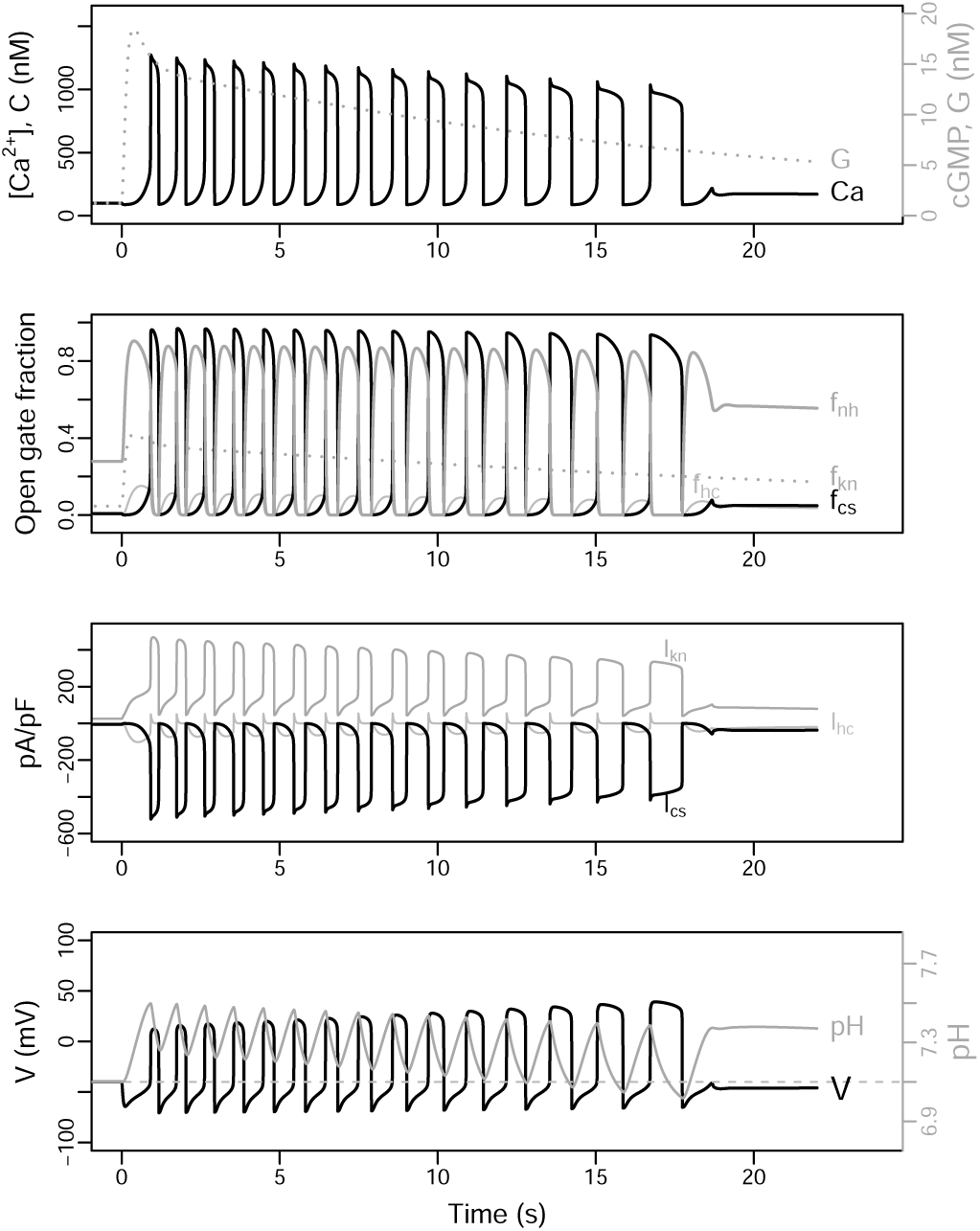
Dynamics of the model featuring the CatSper module (state space 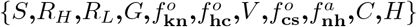). As in Fig. 4, the initial stimulus is *S*(0) = 25 nm, and the graphs are the numerical solutions for the indicated variables and intermediate quantities, obtained with the parameters listed in Tab. A1.

The rise and fall of the cGMP produced by the upstream module leads to a Ca^2+^-spike train with a slow decaying amplitude and an increasing interspike interval (Fig. 6, top). The nadir of the Ca^2+^-oscillations is close or even below the resting level, in contrast with the oscillations predicted by the alternative module (Fig. 4). A similar oscillatory behaviour with similar period but different phases is observed on the fractions of open channels (Fig. 4, second row), the respective currents (third row), as well as the membrane potential and proton concentration (bottom).

Compared to the Ca_V_+BK module, the pattern of the Ca^2+^ oscillations is differently shaped by the CatSper module. The individual Ca^2+^-spikes are more prolonged and the corresponding nadir is relatively shorter. The decay of the spike train envelop is less pronounced for the reference parameters in the CatSper module than in the Ca_V_+BK module, while the increase in the interspike interval is more marked in the former than in the latter. It is remarkable that the CatSper module does not predict the sustained plateau of the Ca^2+^-oscillation nadirs present in the Ca_V_+BK module. Finally, another interesting difference between the two modules is the envelop of the voltage spikes whose amplitudes decrease progressively in the Ca_V_+BK module and increase in the CatSper module (bottom in Figs. 4 and 6).

The basis for the limit cycle with CatSper is qualitatively straightforward. The hyperpolarisation increases the activity of the Na^+^/H^+^ exchanger, this triggers the extrusion of protons and accordingly raises pH_i_. As a consequence of this alkalinisation, the voltage sensitivity of CatSper shifts to lower values (see Fig. S3D) and some of these channels open. The inward Ca^2+^ currents rise the membrane potential which in turn tends to increase the fraction of open channels, in a positive feedback loop. However, the progressive depolarisation of the membrane reduces the activity of the exchanger and the pH_i_ tends to return to its basal level, shifting the CatSper voltage sensitivity back to higher values. This, together with Ca^2+^-inactivation of CatSper, allows the hyperpolarising current to surpass the depolarising current of CatSper (see Fig. 6, third row). The cycle restarts when the membrane becomes again hyperpolarised by the still ongoing KCNG current (Fig. 6, second and third row), leading to a new round of enhanced exchanger activity, a transient raise in pH_i_, and the recovery of CatSper currents that eventually overcome the hyperpolarising currents.

As in the case of the Ca_V_+BK module, the bifurcation analysis of the model featuring the CatSper module but with constant input *G* reveals a rather complex dynamical structure. The system displays a cusp involving two saddle-node bifurcations, which, in contrast to the module Ca_V_ + BK, is in lower *G* ranges, such that two [Ca^2+^]_i_ stable equilibria coexist close to the resting state. At higher cGMP values, there is a regime where the stable equilibrium solution coexists with a stable limit cycle, whose maxima and minima are indicated by a succession of black circles in the top graph in Fig. 7. It is worth noticing that the coexistence of stable attractors within a given range of *G* values suggests the presence of bifurcation points that determine the transition of one behaviour to the other, at the boundaries of that range. However, for this model, with the available bifurcation analysis software (e.g. XPPAUT [25]), it was not possible to discover these points or the limit cycle. In fact, the stable limit cycle was detected by solving the model numerically, exploring the phase space by randomisation of initial conditions of the system and allowing it to evolve for long times (*>*5 × 10^5^ iterations). Also an exploration of initial conditions between the two attractors, suggests that a possible presence of highly interlaced attraction basins, which makes numerical calculations difficult (data not shown).

**Figure 7.**
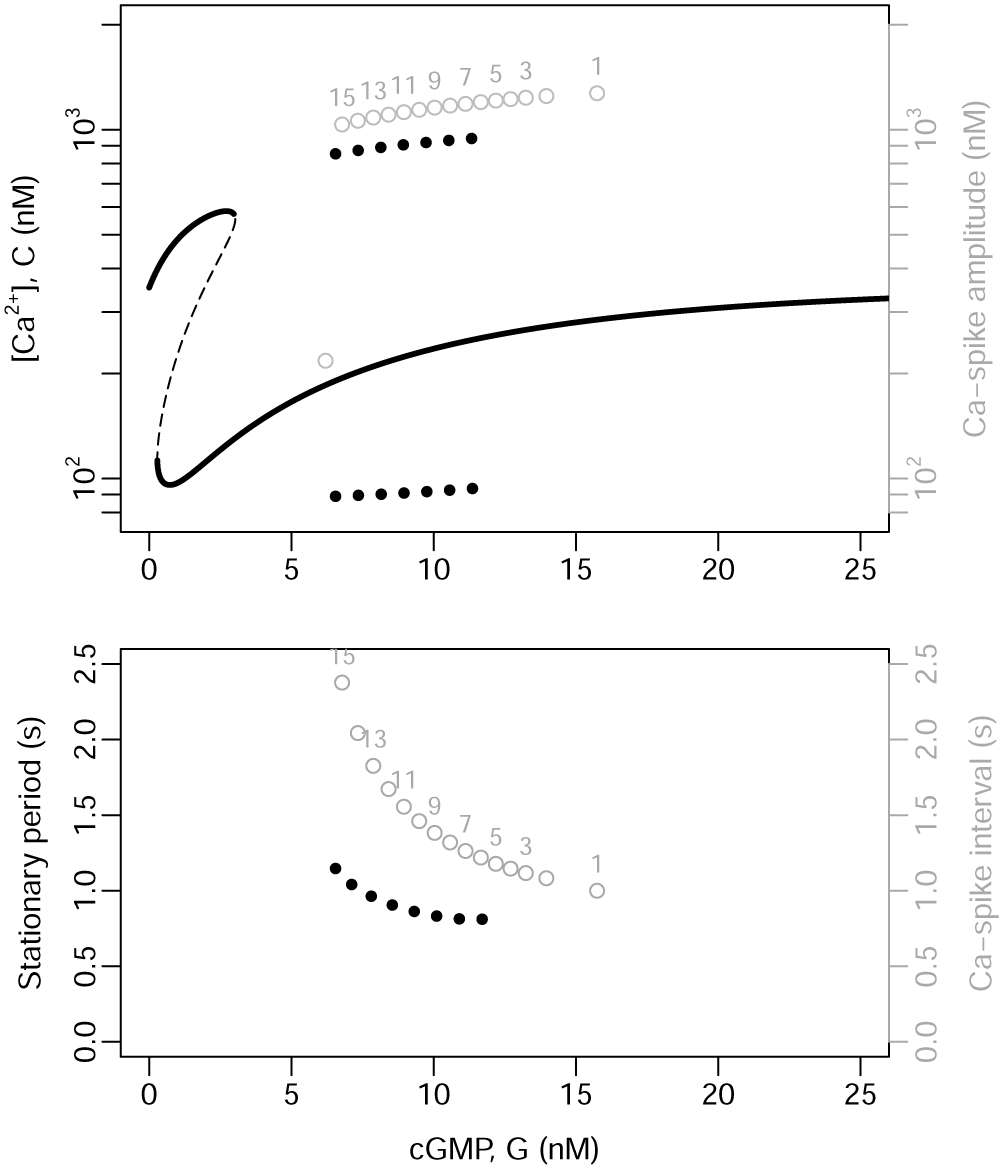
Bifurcation analysis of the model featuring the CatSper module. The analysis was done under conditions wherein cGMP is defined as a constant input (*G* =constant) and upstream variables ignored reducing the state space to 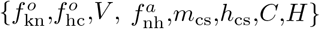. Top: The graphs depict the variable *C* as a function of the input parameter *G*. The thick continuous lines are the stable equilibria and the dashed lines indicate unstable equilibria, as obtained by XPPAUT [25]. The thick dotted lines are the maxima and minima of the stable limit cycle obtained by numerical solutions of the system under random initial conditions. The numbered gray dots indicate the maxima and minima of the consecutive spikes obtained by solving numerically the full model, corresponding to those depicted in Fig. 6. Bottom: The dotted lines represent the period of the stable limit cycles as a function of the constant input *G* obtained by numerical solutions. The numbered gray dots correspond to the interspike interval in the numerical solutions, as in the top graph.

For this scenario, as was the case with the Ca_V_ + BK module, the amplitude decreases and the period of the limit cycle increases (Fig. 7, bottom) as the constant input of cGMP returns to basal levels after having reached high values.

The patterns of the [Ca^2+^]_i_ fluctuations obtained by solving the full model with the CatSper module (Fig. 6) are now amenable to a quantitative interpretation. As the cGMP concentration increases rapidly from the resting state, the system falls within the basin of attraction of the limit cycle determined for constant *G* values; and as the cGMP returns slowly to its basal level, the time required to approach those limit cycles decreases. This can be seen in Fig. 7 by means of the empty circles superimposed on the bifurcation diagram, which show how the system tends towards this limit cycle with a spike amplitude reduction and an increase in the interspike intervals. The differences between the asymptotic values of amplitude and periods predicted by this variant of the model with fixed values of *G* (black circles), and those of the actual peaks obtained when *G* varies in time (empty circles numbered) are more marked in this module than in the case of the Ca_V_+BK module.

### 3.4 Ca_V_+BK and CatSper modules predict distinctive responses to controlled manipulations of intracellular pH and of membrane potential

One of the most direct ways of distinguishing which of the two modules is the most adequate to describe the essential structure of the pathway activated by SAPs is to test their coupling with the pH_i_ dynamics. If sNHE and [H^+^]_i_ are coupled with the Ca_V_ + BK module, the former would be activated by transient hyperpolarisation, and the proton concentration would turn out to be just an output variable, in the sense that pH_i_ does not feedback on any of the processes that determine Ca^2+^ fluctuations; in other words, the concentration of protons does not appear in any of the equations that govern the variables of that model variant. Therefore, this module would predict that changes in the pH_i_ should not entail major changes in the Ca^2+^-spike trains. In contrast, in the CatSper module, the proton dynamics is tightly coupled to the Ca^2+^ oscillations and is an essential part of the mechanisms underlying the oscillatory behaviour. In our analysis of the parameter dependence of the CatSper module, we systematically found oscillations in [Ca^2+^]_i_ that were concomitant with pH_i_ oscillations having the same period, albeit phase differences. Parameters that lead to slower proton dynamics such that the pH_i_ oscillations vanish, resulted in cancellation of the oscillation in [Ca^2+^]_i_. It is worth noticing that attempts to detect periodic pH_i_ oscillations in sea urchin sperm have failed hitherto (see [23]; unpublished observations), which might be explained by experimental limitations or simply because periodic pH_i_ oscillations do not happen *in vivo*. Nevertheless, the different dependence of the two modules on pH_i_ offers a means to distinguish their possible role experimentally.

Suppose that one could artificially raise and fix the pH_i_ inside the flagellum. In this situation, the Ca_V_+BK module would predict that the Ca^2+^-fluctuations would not be affected, while the CatSper module predicts that sustained pH_i_ increase would cancel [Ca^2+^]_i_ fluctuations and this cation would be maintained at higher concentration. This effect is illustrated in Fig. 8A and B, where we also overlay the results of registering Ca^2+^ dynamics in the presence of ammonium chloride (NH_4_Cl), an agent that arguably raises the intracellular pH without affecting any of the other channels modelled here. As observed in the experimental trace, the exogenous alkalinisation gives rise to a dynamics that is qualitatively similar to the one predicted by the pH_i_-dependent CatSper module.

**Figure 8.**
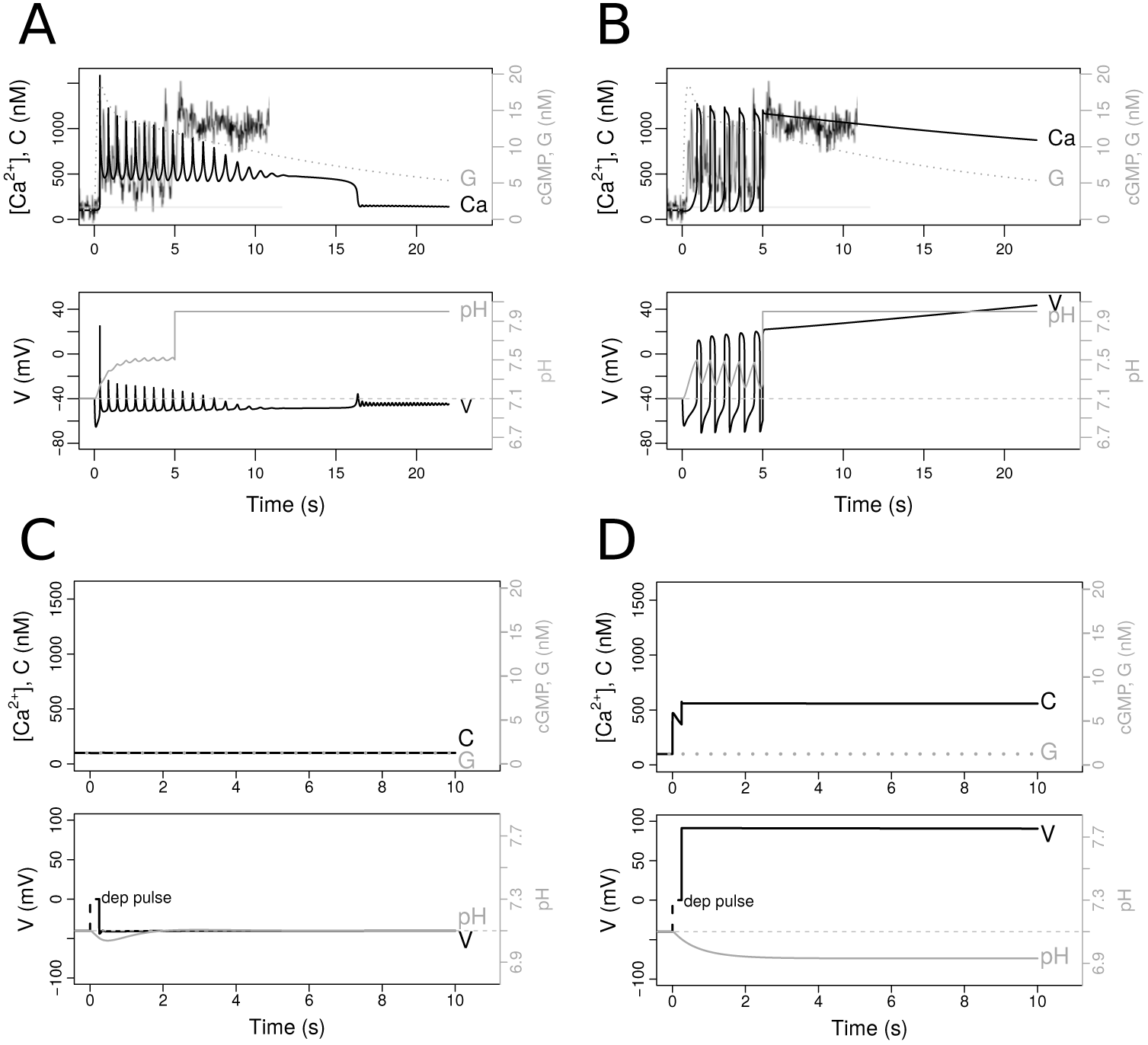
Ca_V_+BK (A,C) and CatSper (B,D) modules predict distinctive responses to manipulation of intraflagellar pH (A,B) and membrane potential (C,D). The graphs show the numerical solutions of the indicated variables under the reference parameters as in Fig. 4 and 6. In A and B, a normal response to SAP develops during the first 5 s, and is then perturbed by an artificial raise in pH_i_ to a constant value maintained thereafter. Variables related to pH_i_ and sNHE are coupled to Ca_V_+BK module with the same reference parameters used in the CatSper module. The noisy gray line is a rescaled trace of the intensity of a pH-sensitive fluorescent probe in *S. purpuratus* sperm cells (obtained as described in [23]). In C and D, the normal resting state is perturbed (in the absence of SAP, *S* =0 nm) by setting the membrane potential to a higher value (*V* =0 mV) at time t = 0 for a short period of 0.25 s. The voltage dynamics during the exogenous stimulus is indicated by dashed lines.

Another distinct feature of the two candidate modules is that at resting state the Ca_V_+BK module has a single stable equilibrium, while the alternative module with CatSper predicts the coexistence of two possible stable equilibria, characterised by [Ca^2+^]_i_ at either basal levels or above basal levels. This is best seen in the bifurcation diagrams in the top graph of Fig. 7. A temporal raise in membrane potential, according to the Ca_V_+BK module, may lead to an increase in [Ca^2+^]_i_ that would rapidly return to its unique steady state (Fig. 8C). In contrast, according to the CatSper module, a sufficiently strong perturbation of the membrane potential may force a switch from the lower basal steady state to the state characterised by higher [Ca^2+^]_i_, where it will remain (Fig. 8D). As a corollary of these properties, the Ca^2+^-spike train will always terminate in the basal level according to the Ca_V_+BK module, while according to the CatSper module, the [Ca^2+^]_i_ may remain at high values after the spike trains vanish under particular dynamics that favour depolarised membrane potential, as is the case of the time course depicted in Fig. 6.

### 3.5 The output of the CatSper module is tuned by the Ca_V_ + BK channels if present in low proportions

In the previous sections, we have shown that both alternative modules, once coupled with the upstream module, can semi-quantitatively describe the observed data. These two modules share some features and potentially could coexist in the flagellum being coupled through the membrane potential, pH_i_, and [Ca^2+^]_i_. To investigate how the coupling of the modules would affect the dynamics, we considered a scenario in which the actual composition of the system would be a weighted combination of the two modules by means of a weight parameter given by *θ*. For *θ* = 0, one recovers the submodel with CatSper module alone, for *θ* = 1 one recovers the submodel featuring Ca_V_+BK module only, and when *θ* = 0.5 the total conductance of the components in each module are half of their respective reference values. To ensure the recovery ot the aforementioned extreme scenarios, in our new weighted scheme, the parameters that regulate the resting membrane potential and ion concentrations were weighted as well (see equation 9).

The importance of how the pH_i_ dynamics influences the [Ca^2+^]_i_ response was illustrated in the previous section, so we now take the CatSper module as our reference scenario since it contains the necessary components to account for this regulatory mechanism. We titrate in the components of the Ca_V_+BK module under three combination schemes. First, we explore the addition of Ca_V_ and BK separately (Fig. 9), then we add both components maintaining their conductance ratio as in the original module Ca_V_+BK (Fig. 10). This simple analysis shows that the modules cannot be combined freely without compromising the dynamics.

**Figure 9.**
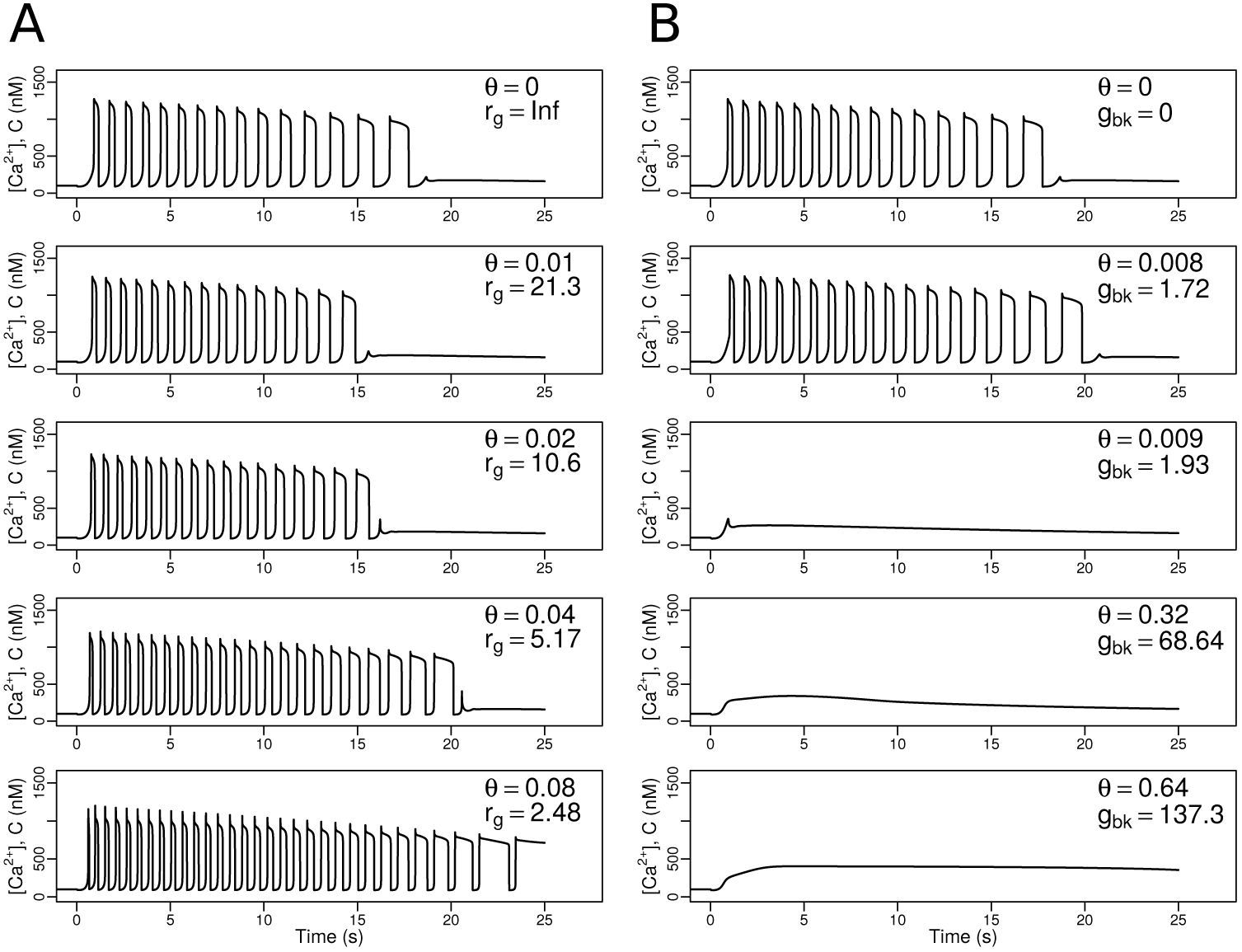
Titration of Ca_V_ or BK channels on the CatSper module. In A and B, we show numerical solutions for calcium under different values of the weighting parameter *θ*. As reference, the original scenario of only CatSper, i.e. *θ* = 0, is shown in the top row, while the subsequent rows correspond to the gradual increase of *θ*. In A, the reference value of the Ca_V_ conductance density, *g*_cv_, is multiplied directly by *θ*, whereas the reference value of CatSper conductance density is multiplied by (1 - *θ*); the resulting ratio of CatSper/Ca_V_ conductance densities, *r_g_*, is shown for each *θ* value. The higher the value of *r_g_*, the greater the predominance of CatSper with respect to Ca_V_. In B, *θ* sets the percentage of the reference value of *g*_bk_, the conductance density of BK; unlike scenario A, the conductance density of CatSper is not weighted by *θ*. For each titration, the corresponding modified *g*_bk_ value is shown

**Figure 10.**
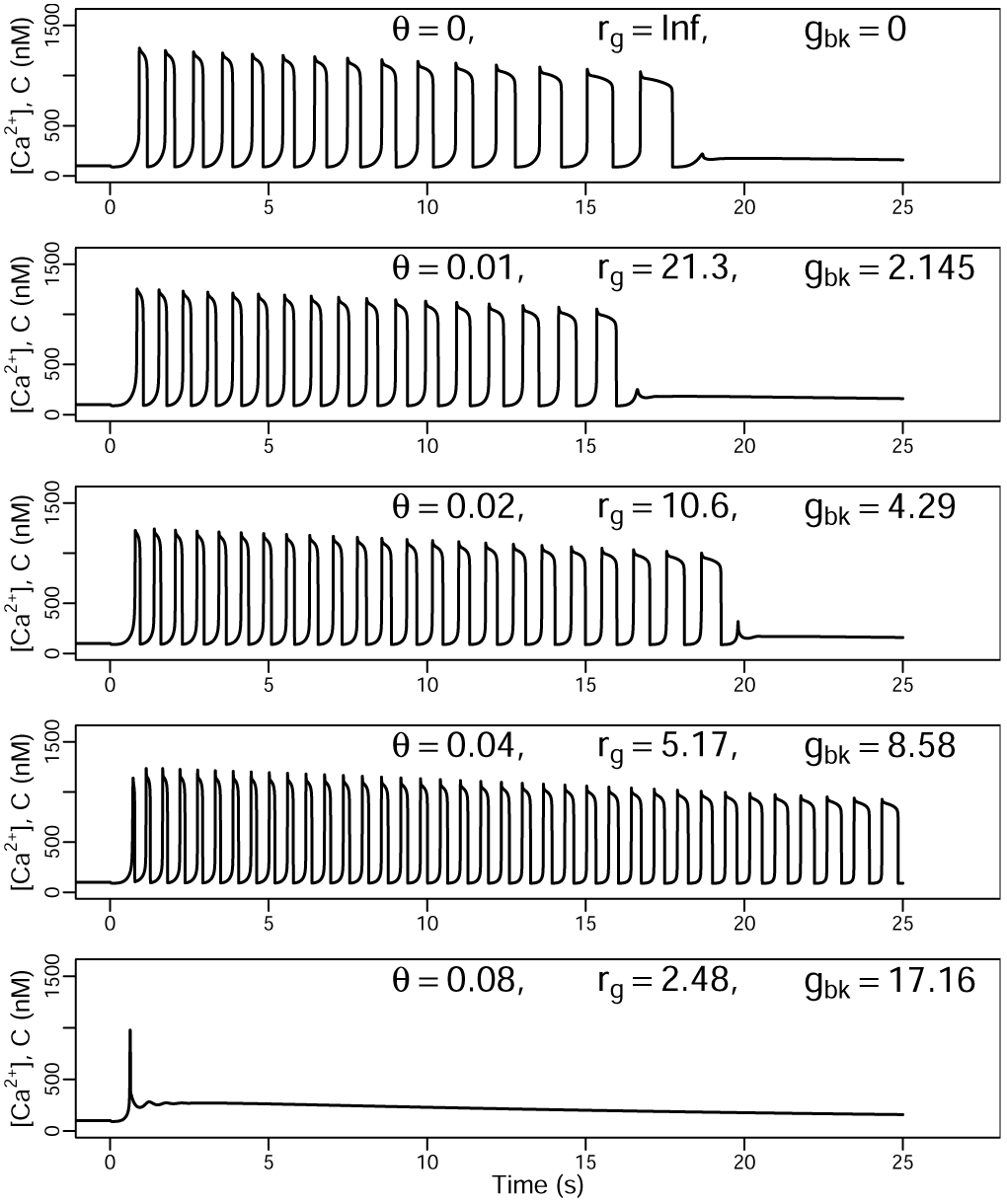
Titration of the module Ca_V_ + BK on the CatSper module. Starting from the model that includes the CatSper module, numerical solutions for calcium are shown under different values of the weighting parameter *θ*, which controls the percentage of module Ca_V_ + BK that is being added. *θ* multiplies the reference conductance *g*_cv_ and *g*_bk_, and correspondingly the conductance density of CatSper decreases by the factor (1 - *θ*). For reference, the original CatSper-only scenario is shown first, i.e. *θ* = 0, and the subsequent rows correspond to the gradual increase of *θ*. The effect of this parameter on the channel densities is reported in the coefficient *r_g_*, which measures the total conductance ratio CatSper/Ca_V_, as well as the new values *g*_bk_. As in the figure 9, the greater the value of *r_g_*, the greater the predominance of CatSper on the total calcium conductance with respect to Ca_V_.

The most noticeable effect of adding relatively small titers of Ca_V_ to the CatSper module, depicted in Fig. 9A, is the overall response acceleration without qualitatively modifying the envelop or the shape of the peaks. For the first two lower titers of Ca_V_ (second and third rows), the frequency increases but the total duration of the spike train is shortened compared to the original CatSper scenario (first row). For the third titer, the response is both accelerated an extended. However, by the fourth titration (last row), the response lasts for tens of seconds and tends to a high calcium level instead of approaching the resting state, these features being incompatible with what is observed experimentally.

The second combination scheme in which BK is added to the CatSper module (Fig. 9B), shows how extremely sensitive this module is when coupled with a channel that has the characteristics of BK. From very small values of conductance density, *g*_bk_, the oscillations disappear while leaving only a tonic calcium increase that reaches intermediate values above the basal level (rows 3 and onwards). Nonetheless, in considerably small values of *θ*, the capacity of the system to oscillate is able to cope with the addition of BK to some extent; this is seen by a change in the time scale of the response without altering the qualitative form of the peaks (second row). In the latter case, there is only a subtle increase in frequency, perceivable by considering that in the first 15 s there are 13 fluctuations without BK, whereas with BK there are 14. Regarding the total total length of the spike train, with BK the response ends just 2 seconds later compared to the CatSper module alone.

When analysing the most complex scenario, consisting of the combination of the two modules, similar effects to the observed in the titration of only Ca_V_ were retrieved, but at different values of *θ* (Fig. 9A), i.e. the increase in the frequency of oscillations without introducing significant changes to the shape of the peaks or to their envelop, as well as the response lengthening. On the other hand, when comparing the two previous schemes with the mixed one, it is suggested that only when BK is added together with Ca_V_, the CatSper-dependent response is more tolerant to the presence of that channel at higher conductance densities (compare values of *g*_bk_ in figures 9B and 10); moreover, BK is also responsible for extending the duration of the spike train (e.g. compare the third row of figures 9A and 10). However, above a certain level of *g*_bk_ (last row), the oscillatory response collapses, leaving only an initial transitory increase; this confirms the negative effect of BK on the spike trains at high densities, as in Fig. 9B.

## 4 Discussion

This article explored the collection of channels that can explain the dynamics of the [Ca^2+^]_i_-spike trains elicited by SAP in sea urchin sperm. The large number of parameters (Table A1) used in the models studied here would deter any attempt of this kind, if it was not for the availability of quantitative time series of cGMP and membrane potential in response to several SAP concentrations. The parameters of the early module that were not obtained from the literature were estimated by fitting the model to these time series data. This was possible given the structure of the signalling network in which the upstream components are independent of the dynamics downstream and the fact that the model predicts a smooth non-oscillatory dynamics. The early signalling module was able to reproduce the temporal evolution of its output variable cGMP and also the early membrane potential changes in response to different SAP concentrations, measured in cell populations [8, 10, 20]. This is remarkable considering that this module makes very simple assumptions about the receptor’s ligand binding and guanylate cyclase activity as well as about cGMP dynamics, neglecting adaptation mechanisms such as sensitivity readjustment of the receptor affinity that have been documented [37–39]. A noteworthy observation is that the cGMP dynamics at longer times, is characterised by a slow decay in which the cGMP concentration is in quasi-steady state following the slow decay of the receptor form with low guanylate cyclase activity (*R_L_*) (Sec. A1.1, Fig. 2, first and second row). If the model predicted that cGMP itself would undergo periodic oscillations (arising potentially from feedbacks of pH_i_ on phosphodiesterase activity [40]), instead of a smooth dynamics, then we would lose the inferential power of fitting the predicted cGMP value to the mean population cGMP levels.

The upstream module alone was able to explain the early hyperpolarization dynamics based on the joint action of KCNG and spHCN channels (Fig. 2, bottom). The parameters were set such that, in the absence of any other channels, the repolarisation initiated by spHCN following the hyperpolarising current of KCNG, leads to a partial recovery of the membrane potential that remains below the resting value. The partial recovery implies that spHCN effective conductance is sufficient to set into motion the downstream modules (e.g. the opening of closed Ca_V_ channels or shifting to higher opening rate of the voltage-gate of CatSper) but not too strong to prevent the oscillatory [Ca^2+^]_i_ dynamics (results not shown). This property of the signalling network is amenable to experimental corroboration by assessing whether the membrane potential recovery is only partial under a setting in which all calcium entry is prevented by using zero external concentrations of this cation.

The key question was what is the composition of the downstream module that links the SAP-induced cGMP response to the downstream [Ca^2+^]_i_ fluctuations. Our analysis indicates that either one of two modules featuring voltage-dependent channels could be responsible for the dynamics of [Ca^2+^]_i_ fluctuations. The first module calls into action Ca_V_ channels that undergo voltage-dependent sequential transitions from inactivated to closed, to open and back to inactivated states. These irreversible transitions bring forth a limit cycle with constant period with an amplitude that is controlled by cGMP activity. These Ca_V_ channels, which are predominantly inactivated at the resting potential, are released from inactivation upon a transient hyperpolarisation and subsequently opened when spHCN channels activity raises the membrane potential. In this module, the oscillations are strongly modulated in amplitude and tempo by the Ca^2+^-dependent channel BK to produce the experimentally observed progressive increase in the intervals between [Ca^2+^]_i_ spikes. The oscillations predicted by this module vanish when cGMP falls below a critical value involving a Hopf-type bifurcation (Fig. 5A).

The second module capable of explaining the organisation of the [Ca^2+^]_i_ fluctuations features a single channel, CatSper, that is voltage- and pH-dependent and is reversibly inactivated by intracellular Ca^2+^ ions. In contrast with the previous module, which is set in motion directly by the effect of transient hyperpolarisation that relieves Ca_V_ channels from inactivation, the initial opening of CatSper channels is triggered indirectly by alkalinisation of the cytoplasm driven by the hyperpolarisation-dependent Na^+^/H^+^-exchanger activity (Fig. S3). The cycling behaviour is produced by the negative feedback that depolarisation driven by CatSper activity has on inactivating the Na^+^/H^+^-exchanger, reducing pH_i_ and decreasing CatSper activity. Another feedback arising from the inactivation of CatSper by [Ca^2+^]_i_ plays a more preponderant role in controlling the length of the Ca^2+^-spike train and in increasing the interspike intervals. As in the case of the first module, the oscillations generated by the module stop when the cGMP activity falls below a critical value (Fig. 7).

It is noteworthy that, under the reference parameters used in this article, the overall temporal scale of the calcium response produced by the CatSper module is slower compared to the experimental tracings, in terms of real time units; however, if the relative progressive increase of interspike intervals is contrasted instead, this property becomes more marked in this model than in the one featuring the Ca_V_ + BK module, and more closely related to the experimentally observed (Fig. 3C). In the numerical solutions with the former module the interval between the last spikes can be two or three fold larger than the interval between the first spikes in the train, as observed in the experiments in single cells. A comparable increase in interspike intervals can be obtained with the Ca_V_+BK module under different parameter regimes (not shown). Conversely, the progressive decrease in the amplitude observed experimentally is better accounted for by the Ca_V_+BK module than by the CatSper module under the reference parameters used here. A more evident decline of the envelop of the Ca^2+^-spikes can be obtained using the CatSper module under other parameter regimes in which the spikes became less realistically interspersed in time.

These two downstream modules confer some distinguishable properties to the signalling network when coupled to the upstream module. One of those properties is the envelop of the membrane potential oscillations. In the module featuring the Ca_V_ + BK channels pair. This envelop has a shape similar to that of the [Ca^2+^]_i_ spikes. In contrast, the CatSper module predicts that the amplitude in the membrane potential peaks should increase until it suddenly ceases (Fig. 6, bottom), thus establishing an opposite trend to the calcium-spike train envelop. Interestingly, the ascending *V* envelop is more similar to the observed at prolonged times in membrane potential measurements of sperm populations stimulated with high doses of SAP [8].

Another good example of distinctive features between the alternative scenarios is the permissive *G* range that allows for oscillatory solutions. At too low (close to resting) or too high intracellular GMP concentrations, the model with CatSper predicts a sustained rise [Ca^2+^]_i_ without noticeable oscillations (Fig. 7). Conversely, in the Ca_V_ + BK module, the periodic regime is maintained for any value above a critical level of cGMP, however, above more elevated concentrations, the amplitude and period saturate to nearly constant values (Fig. 5A). These properties of the model could be addressed experimentally by phosphodiesterase inhibitors or by elevating cGMP to saturating concentrations by uncaging an analogue. It is worth recalling the reports that intracellular uncaging of cGMP leads to an increase in [Ca^2+^]_i_ that is oscillatory in the case of *A. punctulata* [24, 41] but transient in the case of *S. purpuratus* ([11]) spermatozoa. The discrepancy between the two species could be reconciled if there are species-specific differences in the boundaries of cGMP intervals that produce oscillations or alternatively by experimental differences in the uncaged concentration of the cyclic nucleotide.

Albeit these two candidate modules predict qualitatively the observed calcium responses under particular parameters regimes, we found that the three channels, Ca_V_ + BK on the one hand and CatSper on the other, interfere with each other leading to unrealistic behaviours when the effective calcium currents are of the same order of magnitude (bottom in Fig. 9 and 10). This modelling result suggested that the two modules could be mutually exclusive raising the question of which one is predominating in sea urchin spermatozoa. The present analysis indicates that the two modules predict very distinct responses to the artificial induction of sustained alkalinisation. Hence, the CatSper module, but not the one featuring Ca_V_ and BK channels, predicts that transient or sustained alkalinisation should result in sustained [Ca^2+^]_i_ as observed experimentally following the addition of NH_4_Cl to *S. purpuratus* spermatozoa (Fig. 8A). The confrontation of these two modules with the experimental measurements suggests that the channel responsible for the [Ca^2+^]_i_ responses to SAP is the pH-dependent CatSper instead of Ca_V_.

Further considerations on the above matter are called for. It is worth emphasizing some additional implications of the CatSper module, as these are also amenable to experimentation. Under the reference parameters, the oscillatory regime in the CatSper requires oscillations of proton fluxes and intracellular pH. Rather than inactivation of CatSper by [Ca^2+^]_i_, the effective feedback loop explaining the intracellular Ca^2+^ oscillations is the retroinhibition of the sNHE activity by depolarising currents. In other words, observing [Ca^2+^]_i_ oscillations dependent on CatSper implies necessarily the concomitant observations on pH_i_ oscillations with the same period and different phase. Oscillations of pH_i_ with the same period of those of [Ca^2+^]_i_ were not detected in experiments where the levels of pH_i_ have been monitored in individual cells ([23]; unpublished observations). This might reflect limitations of the experimental system (e.g. too low signal to noise ratio of the pH probe as compared to the Ca^2+^ probe) or, alternatively, it might mean that intraflagellar pH is not oscillating periodically. In the latter case, the hypothesis that CatSper is the main responsible for fluctuating Ca^2+^ currents, at least in the way we modelled it here, is questionable and must be reexamined. In this context, it might be relevant to note that oscillatory fluxes of Ca^2+^ and protons have been reported with the same period as growth oscillations of pollen tubes (reviewed in [42]).

The observation that a rise in pH_i_ precedes the first Ca^2+^ spike in sea urchin sperm stimulated by SAP has been interpreted as an indication that Ca^2+^ currents are a consequence of this early alkalinisation [14, 22, 23]. This interpretation has become a major cornerstone of the hypothesis that pH-sensitive CatSper channel is responsible for Ca^2+^ fluctuations. It is worth calling attention to the early time courses of pH_i_ and [Ca^2+^]_i_ obtained by the numerical solution of the model featuring the Ca_V_+BK module (Fig. 8A). In this model, where Ca^2+^ currents are, by construction, independent and unaffected by pH_i_ one can nevertheless observe an onset of alkalinisation that precedes the first [Ca^2+^]_i_ spike. This illustrates the frailty of inferring causation from time series data in general and raises a cautionary note on the interpretation that a pH-sensitive Ca^2+^ channel must be involved in sea urchin sperm responses to SAP.

Another testable prediction of the CatSper module is the existence of bistability at the resting values of cGMP. This implies that perturbations exist that may lead to sustained [Ca^2+^]_i_ value above the normal resting one, in the complete absence of SAP signals. We illustrated this by a transient rise in membrane potential that locks the system in the stable state characterised by higher, tonic-like [Ca^2+^]_i_ (Fig. 8D). Another perturbation with comparable effects would be a transient yet complete depletion of cGMP that would push the system across a bifurcation point in which the lower equilibrium vanishes. Following such perturbation, hysteresis is predicted such that the system could remain in the higher [Ca^2+^]_i_ state, after restoration of the basal cGMP levels. The CatSper module bistability also implies that after a [Ca^2+^]_i_-spike train terminates, the system may or may not return to the basal level. It is interesting to note that this alternative [Ca^2+^]_i_ equilibrium state also entails low pH_i_ levels and depolarised membrane potential (Fig. 8D, bottom). If such a state existed, its physiological meaning would be associated with a state of quiescence, since the acidification of the cytoplasm below the basal pH_i_ inhibits both the metabolism and the dyneins, which are the molecular motors that drive the flagellar beating [33, 43]. On the other hand, the viability of sperm would be affected in this state, since sustained high levels of calcium represent a universal signal that triggers cell death in the majority of eukaryotic cells [44–46]. These properties of bistability and hysteresis at *G* levels close to the resting state are not present in the Ca_V_+BK module, thus offering yet another way of teasing apart the two hypotheses.

The two modules are almost mutually exclusive. The Ca_V_ and BK channels can fine-tune the temporal structure of spike trains elicited by CatSper if present at minute densities. At densities or currents comparable to those of CatSper these channels will disrupt the [Ca^2+^]_i_ oscillation train. This suggests that the stoichiometry of the channels in sea urchin sperm might be tightly regulated. At least this would be the case, according to the general conviction that the temporal organisation of the [Ca^2+^]_i_-spike trains is critical for steering the spermatozoon to the egg. Such constraints on stoichiometry are difficult to reconcile with the pervasive observation that the number of molecules per cell varies in time and across a cell population. This has been observed in free living cells (eg. [47, 48]) or in cells of multicellular organisms (eg. [49, 50]). If sperm cells require tight control of channel stoichiometry as implied by the results with CatSper, one would expect that the coefficient of variation in the number of channels per sperm cell should be on average lower than the coefficient of variation in membrane receptors of somatic cells, such as haematopoietic or epithelial cells.

In the same vein, the CatSper module predicts [Ca^2+^]_i_-spike trains in agreement with the experimental observations only if BK channel densities are negligible. If one assumes that the unitary conductance of BK channels in marine invertebrates is closer to the estimated in mollusk and crustacean cells [51–53] (under asymmetric conditions that approach physiological gradients) than its counterpart in mammalian cells [54], this would be of the order of ≈70 pS. Under these conditions, the BK conductance densities that allow for oscillatory trains (Fig. 9B, second row) would imply channel counts of less than 1 molecule per flagellum, given the membrane area (Tab. A1). Therefore, although BK is expected to play a key role in a scenario wherein Ca_V_ channels are the ones driving calcium, the role of this channel, if any, should be minimal if CatSper turns out to be the predominant channel. The role for BK in [Ca^2+^]_i_ responses to SAP was inferred indirectly by the alterations of the period of these oscillations by blockers such as niflumic acid [55, 56] and Iberiotoxin [13], which were corroborated by simulations of a discrete model. In the present quantitative framework, there are shared effects of BK both on the Ca_V_+BK and CatSper modules, such as defining a tonic-like minimal value for the nadir of [Ca^2+^]_i_ oscillations and increasing the period of the oscillations. This notwithstanding, the CatSper module, under the reference parameters, precludes the presence of BK in sea urchin sperm at significant levels.

We used bifurcation analysis of a subset of the differential equations of the full system as a heuristic tool to grasp how the temporal evolution of cGMP activity affects the envelop and intervals between [Ca^2+^]_i_ spikes. It is important to bear in mind that the present analysis involves an asymptotic approximation, i.e. in our case *G* was set to a fixed value as a way to transform this variable into a parameter and quantify the amplitude and period of the stable limit cycles. The experimental corroboration of such analysis would require development of a cGMP-clamping technique, analogously to classic voltage-clamping experiments or synthetic biology approaches that seem presently unfeasible in sea urchin sperm.

We end by noting that the present work illustrates the added value of quantitative analysis, obtained by differential equation systems and bifurcation analysis, which beyond offering a framework to characterise mechanisms overcoming experimental limitations, helps to identify the main elements controlling the system. Quantitative modelling allows to define guidelines for future experiments to assess the theoretical predictions, to identify key properties and testable quantities, and to prioritise these properties and quantities according to their impact and significance for the system.

## 5 Author contributions

A.D., J.C., D.A.P.-E. and G.M.-M., devised the research, followed its development, interpreted and discussed the findings. J.C. and D.A.P.-E. performed all the calculations and wrote the manuscript. A.D. provided the experimental content and contributed to the writing and revision of the manuscript. G.M.-M. provided formal mathematical content and contributed to the writing and revision of the manuscript. A.G. performed experimental measurements for Fig. 3. A.L.G.-C. performed experiments of ammonium stimulation for Figs. 8A,B. T.N. provided biological feedback.

## 6 Acknowledgments

Funding: Dirección General de Asuntos del Personal Académico of the Universidad Nacional Autónoma de México(DGAPA-UNAM)-PAPIIT grants IN112514 to G.M.-M. and IN205516 to A.D. Consejo Nacional de Ciencia y Tecnología from Mexico (CONACyT) grants CB-2015-01 255914-F7 to G.M.-M., Fronteras 71 39908-Q to A.D. G.M.-M. thanks PASPA/DGAPA/UNAM for support during a sabbatical leave at the Ecole Normale Supérieure, Paris. J.C. is funded by the Fundaça ̃ o para a Ciência e Tecnologia and Instituto Gulbenkian de Ciência. Daniel Alejandro Priego-Espinosa was a doctoral student from Programa de Doctorado en Ciencias Biomédicas, UNAM and received fellowships 275795 and CB-2015-01 255914-F7 from CONACyT; this work is part of his PhD Thesis. He also thanks UNAM for the fellowships given by Programa de Apoyo a los Estudios de Posgrado (PAEP) and Movilidad Internacional de la Coordinación de Estudios de Posgrado (CEP). Ana Laura González-Cota thanks CONACyT for the postdoctoral fellowship EPE-2017 291231. We especially thank Andreas Bohn and Jesús Espinal for fruitful discussions. The authors declare that they have no competing interests.

## Appendices

### A1 Signalling components and channels

In the following section, we will present the molecular components and channels that were implicated in SAP signal transduction and that are relevant for our study. For each of these components we will describe: i) the evidence for their role in SAP-signalling, ii) a mathematical model for its dynamics, iii) the reference parameter values and how they were obtained.

#### A1.1 SAP-receptor and guanylate cyclase

The binding kinetics of SAP to its respective sperm receptor has been studied in sea urchin species such *L. pictus* [57] and *S. purpuratus* [58], whose spermatozoa respond to the decapeptide speract. The reported values for the association and dissociation rate constants for speract imply that the formation of a ligand-receptor complex is essentially irreversible during a chemotactic response, with an average life time of several hours. Since the time scale of the SAP response is of the order of seconds, for the sake of simplicity we neglected the dissociation rate. In order to explain the kinetic features of experimental cGMP data [20] we propose the following mechanism: once SAP molecules (*S*) bind to free receptors (*R*), they go through several states in an irreversible manner, each with decreasing guanylate cyclase activity, namely, *R_H_* (high), *R_L_* (low) and *R_I_* (inactive). In the context of previous work, *R_H_* and *R_L_* might be considered as equivalent to the phosphorylated and dephosphorylated forms of guanylate cyclase, respectively [38, 59–61]. Receptor activation is modelled by the following kinetic diagram:

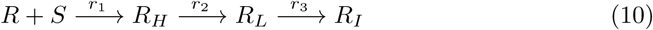

subject to the mass conservation relations:

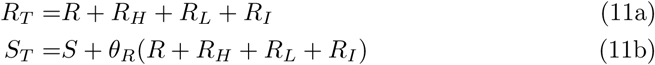

with *R_T_* and *S_T_* being constant values that denote total quantities of receptors and SAP, respectively. The density of the receptor forms (*R_T_*, *R*, *R_H_*, *R_L_*, *R_I_*) are given in units of molecules per cell, and the concentrations of SAP (*S_T_*, *S*) are measured in moles per unit volume, which are brought to the same dimensions by the conversion factor *θ_R_*. When comparing the model to quantitative measurements in bulk we have *θ_R_* = *s/N_A_*, where *s* is the concentration of sperm cells per litre and *N_A_* is Avogadro constant. The dynamics of these receptors forms are described by the following ordinary differential equations:

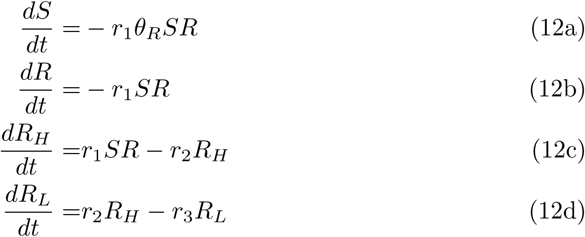

Regarding the activation of guanylate cyclases stimulated by SAP, minor differences exist at the molecular level between sea urchin taxonomic orders. In sperm of *S. purpuratus* and *L. pictus*, which belong to the Echinoida order, receptor and guanylate cyclase have been reported to be two separate yet presumably tightly coupled membrane-bound proteins; once a SAP molecule binds to the former [62] this in turn stimulates the activity of the latter [63]. In the third species, which belongs to Arbacioida order, a single protein has the activities of ligand binding and guanylate cyclase [64, 65]. To capture the commonalities in a single model, we only take the latter and simpler case, thus ignoring any possible delay between SAP binding and guanylate cyclase activation. Thus, the term corresponding to the SAP-dependent cGMP synthesis, *σ_G_*, in equation 4 is defined as:

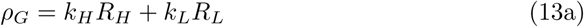

where the constants *k_H_* and *k_L_* measure the guanylate cyclase activities of the high (*R_H_*) and low activity (*R_L_*) SAP-receptor forms, respectively.

#### A1.2 Cyclic nucleotide gated K^+^ channel

The K^+^-selective cyclic nucleotide gated (KCNG) channel is a pseudotetrameric channel with up to 4 cyclic nucleotide binding domains [21, 31], nonetheless, the binding of only a single cGMP molecule is necessary and sufficient to open a channel, thus giving a non-cooperative gating unlike most cyclic nucleotide gated channels [21, 66]. We therefore assume that the fraction of open channels, denoted as 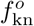, increases with a rate directly proportional to the cGMP activity (*G*) and decreases with first order kinetics.

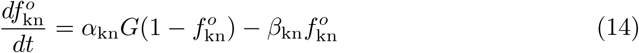

Expression for both the steady-state fraction of open channels, 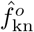, and the characteristic time, *τ*_kn_, can be derived from the opening and closing rates:

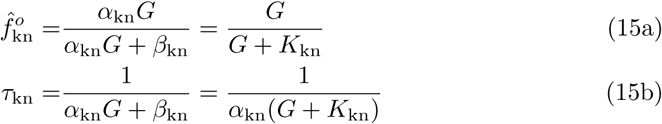

with 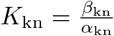. The ion current density through these channels is modelled as:

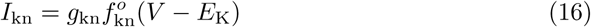

where *E_K_* is the Nernst potential of the K^+^ and *g*_kn_ denotes the effective conductance of the KCNG channel (the product of the unitary conductance by the density of channels).

#### A1.3 Hyperpolarisation-activated cyclic nucleotide-gated channels

Hyperpolarisation-activated cyclic nucleotide-gated channels from sea urchin sperm (spHCN) are cationic channels, weakly selective and carry a mixed current of Na^+^ and K^+^ with a permeability ratio *P*_K_*/P*_Na_ of ~4-5 and a reversal potential of *E*_hc_ =*−*30 mV-*−*10 mV which would allow an inward Na^+^-current under physiological conditions that, when open, depolarises the cell membrane [27–29]. Unlike typical spHCN channels, spHCN-current inactivates in the absence of cAMP, and the latter increases the maximal open-probability instead of shifting the half-activation voltage of the G/V curve [27, 67]. Notwithstanding that, for the ionic current equation associated with this channel, we assumed in the model that cAMP is always present at sufficient concentration to avoid current inactivation, hence any role of cAMP in the overall dynamics is neglected. Following a classical Hodgkin-Huxley type model, we propose that spHCN opening depends on three independent gating variables, denoted by *m*_hc_, such that the fraction of open channels is 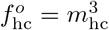. The dynamics of the gating variables is described by:

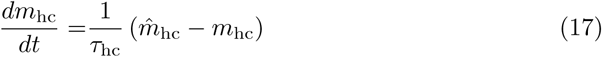

wherein the steady-state fraction of active gating variable, *m ̂* _hc_, is modelled with a Boltzmann equation, whereas the characteristic time, *τ*_hc_, is modelled as a Gaussian function of the voltage:

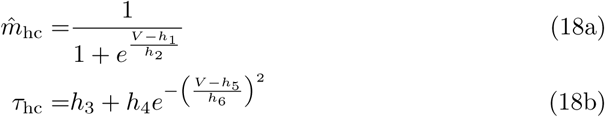

The current via spHCN channels *I*_hc_ is given by:

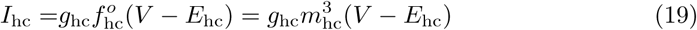

#### A1.4 Electroneutral sodium / proton exchanger

A sperm-specific electroneutral sodium/proton exchanger (sNHE) is expressed in sperm flagella membrane [68]. Its activity is enhanced with hyperpolarised membrane potentials and inhibited with depolarisation. When active, it makes use of the Na^+^ concentration gradient between external and internal media to extrude protons by means of a Na^+^/H^+^ electroneutral exchange mechanism with a 1:1 stoichiometry, consequently increasing pH_i_ [69]. Even though sNHE is not a channel, it is distinguished from the rest of NHE family members in having a voltage sensor domain that is homologous to that of voltage-gated ion channels [9], which might explain the observed voltage-dependent Na^+^/H^+^ exchange in sea urchin sperm in spite of being an electroneutral net exchange [69–72]. We therefore model this voltage dependency as a regular gating mechanism similar to that of Hodgkin-Huxley like equations. The fraction of active sNHE molecules, denoted 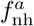, is described by:

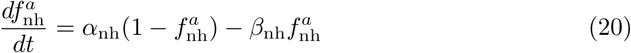

where the activation and inactivation rates are the following functions of the membrane potential:

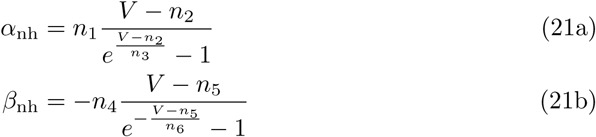

Proton flux as a consequence of exchange activity of sNHE, designated *J_H_* (mol s^*−*1^), is modelled with reversible, rapid-equilibrium, random order bi-bi kinetics[73]:

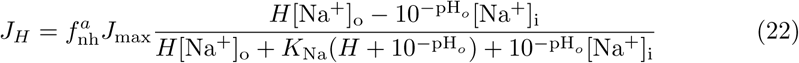

where: *J*_max_ (mol s^*−*1^) is the maximal exchange rate; [Na^+^]_o_ and [Na^+^]_i_ are the extracellular and the intraflagellar concentrations of Na^+^, respectively; pH_o_ is the extracellular pH; and *K*_Na_ is the dissociation equilibrium constant for Na^+^.

#### A1.5 Voltage-gated calcium channels

There has been suggestive evidence for the involvement of classical voltage-gated calcium channels in the SAP-activated response, i.e. Low-Voltage-Activated [8, 10] and High-Voltage-Activated channels [12]. Here we focus on T-type calcium channels, which belong to the first class and are known by being associated to spiking activity in other cell types. We model their kinetics by the following diagram, in which illustrates that they transit irreversibly through three states, namely, inactive Ca_V_^i^, closed Ca_V_^c^ and open Ca_V_^o^:

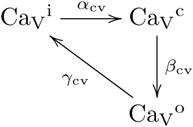

The gating rates are themselves voltage-dependent exponential functions similar to those of Hodgkin-Huxley (HH) type models:

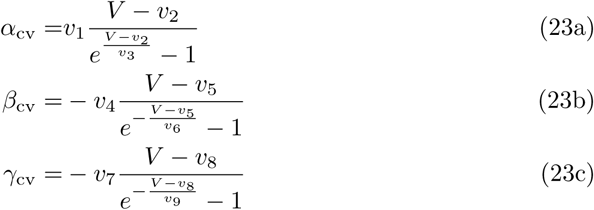

Each transition rate is promoted at a different voltage range: recovery from inactivation (i to c) is favoured under hyperpolarisation (under the reference parameters we have *v*_2_ =*−*55 mV), inactivation (o to i) takes place under marked depolarisation (*v*_8_ =*−*18 mV) and opening of closed channels (c to o) is favoured from potentials slightly hyperpolarised (*v*_5_ =*−*39 mV).

Denoting the fraction of open, closed and inactive channels as 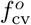, 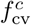 and 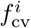, respectively, we have two differential equations for the kinetics of the first two states, whereas the last one is calculated from the conservation equation 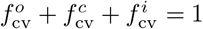.

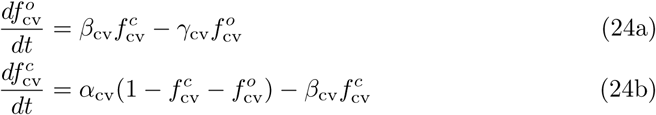

After solving gating equations for equilibrium, we get:

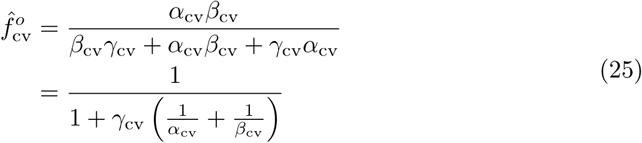

The ion current density through these channels is modelled as:

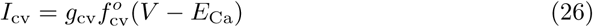

#### A1.6 CatSper channels

The sea urchin genome contains the gene set necessary for the expression of the four core and the three auxiliary subunits that constitute the CatSper channel[74].

Furthermore, the expression of these subunits has been demonstrated in *A. punctulata* sperm flagella [14] and most of them in *S. purpuratus* [15]. In order to model CatSper gating, we consider three regulatory mechanisms reported in studies from mouse, human and sea urchin sperm: voltage-dependence, modulation of voltage-sensitivity by pH_i_, and inactivation by intracellular Ca^2+^ [14, 75–77]. We implement these mechanisms using two independent gating variables, *m*_cs_ and *h*_cs_, representing the fraction of channels with their gate modulated by both voltage and pH_i_ open, and the fraction of channels with Ca^2+^-dependent gate that is not inactivated by calcium. The fraction of open CatSper channels conducting Ca^2+^ currents, 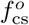, is given by the product of these gating variables:

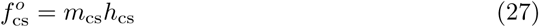

whose dynamics is described by the following differential equations:

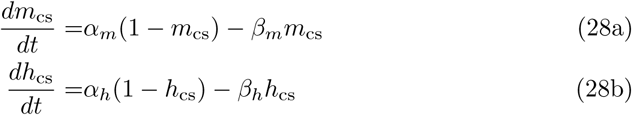

The opening and closing rates of the voltage- and pH-dependent gate were chosen to be the following symmetric functions:

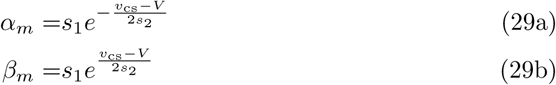

such that the equilibrium open gate curve is a Boltzmann function of the voltage:

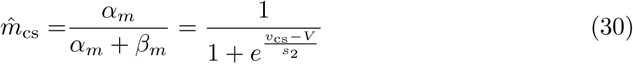

The dependence on pH is ensured by defining the half-maximal voltage, *v*_cs_, as the following function of the proton concentration *H*:

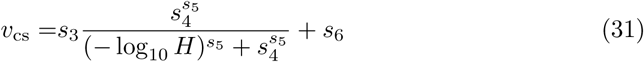

in order to produce a shift of the gating curve towards more negative membrane potentials upon alkalinisation of cytosol, as observed in [14, 75, 76].

The second gate, inhibited by [Ca^2+^]_i_, is assumed to switch between open and closes states with the following rates:

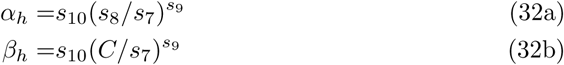

whose function forms were chosen such that when solving for the open gate fraction at equilibrium, *h ̂* _cs_, one obtains a Hill-function:

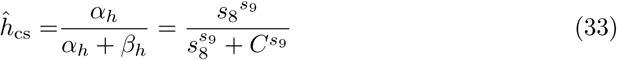

The ion current density through CatSper is modelled as:

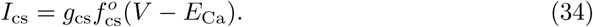

#### A1.7 Big-conductance potassium channels

Big-conductance potassium channels, known as BK (Slo1), are activated by [Ca^2+^]_i_. These channels have been implicated in sea urchin sperm signalling by pharmacological blocking assays [13, 55] and adopted in discrete models [13, 15, 17]. For their dynamics we assume two states, open or closed, where the open-channel fraction, denoted by 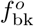, is described by the following differential equation:

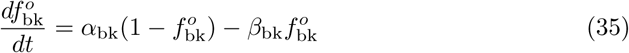

with the closing rate kept as a constant *β*_bk_ and the opening rate being the following Ca^2+^-dependent function :

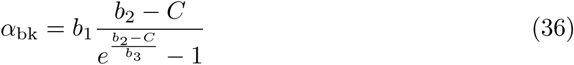

The ion current density through these channels is expressed as:

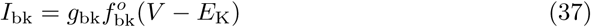

### A2 Variables, intermediate functions and parameter values

Table A1 systematically gathers all the information regarding the quantities used in the model. According to the nature of each of these, the fourth column specifies a numerical value if it is a parameter, or the respective label whether it is a variable or a function. The last column may contain:

- a bibliographic reference if values were used as reported in the literature
- the label F if it is a parameter fitted with functions of the software specified in Materials and Methods section, as well as the bibliographic reference from which the experimental data were extracted to do fitting
- the label MF if it is a parameter that was manually fitted, and the bibliographic reference if any experimental observations were used as guidance
- the label D, if it is a parameter derived by means of an equation, accompanied by the reference to the section where its derivation is explained
- a numeric reference to the corresponding equation when it is a variable or a function.

**Table A1.** Variables, intermediate functions, and parameters values. F=Data-fitted parameter, MF=Manually fitted parameter, D=Derived parameter

| Symbol | Description | Units | Reference value | Reference |
| --- | --- | --- | --- | --- |
| **Upstream module: SAP Receptor and cGMP signalling** |  |  |  |  |
| SAP and receptor |  |  |  |  |
| $S$ | Extracellular SAP concentration | nM | Variable | eq. 12 |
| $R$ | Number of free SAP receptors per flagellum | | Variable | eq. 12 |
| $R_H$ | Number of high activity SAP-receptor complexes per flagellum | | Variable | eq. 12 |
| $R_L$ | Number of low activity SAP-receptor complexes per flagellum | | Variable | eq. 12 |
| $R_T$ | Total number of SAP receptors per flagellum | | $7.36 \times 10^5$ | F [20], Fig. S1 |
| $\theta_R$ | Receptor number to concentration conversion factor | nM | $6.14 \times 10^{-5}$ | D (sec. A1.1) |
| $s$ | Sperm density in measurement chamber | $\text{mL}^{-1}$ | $3.70 \times 10^7$ | F [20], Fig. S1 |
| $r_1$ | SAP-receptor association rate constant | $\text{nM}^{-1} \text{s}^{-1}$ | $2.70 \times 10^{-2}$ | [58] |
| $r_2$ | $R_H$ to $R_L$ conversion rate constant | $\text{s}^{-1}$ | 8.70 | F [20], Fig. S1 |
| $r_3$ | $R_L$ Inactivation rate constant | $\text{s}^{-1}$ | $5.70 \times 10^{-2}$ | F [20], Fig. S1 |
| cGMP |  |  |  |  |
| $G$ | Intraflagellar cGMP concentration | nM | Variable | eq. 4 |
| $G_r$ | Resting cGMP concentration | nM | 1.25 | F [20] |
| $\sigma_G$ | Receptor-independent cGMP synthesis rate | $\text{nM s}^{-1}$ | 23.65 | F [20], Fig. S1 |
| $\delta_G$ | cGMP turnover rate constant | $\text{s}^{-1}$ | 18.92 | F [20], Fig. S1 |
| $\rho_G$ | Receptor-dependent cGMP rate | $\text{s}^{-1}$ | Function | eq. 13 |
| $k_L$ | cGMP synthesis rate constant per $R_L$ | $\text{s}^{-1}$ | 3.25 | F [20], Fig. S1 |
| $k_H$ | cGMP synthesis rate constant per $R_H$ | $\text{s}^{-1}$ | 40.20 | F [20], Fig. S1 |
| $B_G$ | Flagellum cGMP buffering power | | $5.49 \times 10^3$ | F [8] |
| $\theta_G$ | Conversion factor to turn cGMP molecule number into effective intraflagellar concentration | nM | $1.89 \times 10^{-4}$ | D (sec. 2.1.2) |
| KCNG channels |  |  |  |  |
| $f_{\text{kn}}^o$ | Fraction of open channels | | Variable | eq. 14 |
| $\alpha_{\text{kn}}$ | Opening rate constant | $\text{nM}^{-1} \text{s}^{-1}$ | 10 | F [8] |
| $\beta_{\text{kn}}$ | Closing rate constant | $\text{s}^{-1}$ | 257 | [21] |
| $g_{\text{kn}}$ | Conductance density | $\text{pS } \mu\text{m}^{-2}$ | 135.30 | A<sup>a</sup> [8] |
| $I_{\text{kn}}$ | Current density | $\text{fA } \mu\text{m}^{-2}$ | Function | eq. 16 |

Table A1 – continued from previous page
| Symbol | Description | Units | Reference value | Reference |
| --- | --- | --- | --- | --- |
| spHCN channels |  |  |  |  |
| $m_{\text{hc}}$ | Fraction of open gates | | Variable | eq. 17 |
| $f_{\text{hc}}^o$ | Fraction of open channels | | Function | $m_{\text{hc}}^3$ |
| $\hat{m}_{\text{hc}}$ | Steady state fraction of open channels | | Function | eq. 18 |
| $\tau_{\text{hc}}$ | Characteristic time of gating | s | Function | eq. 18 |
| $h_1$ | Half-activation voltage | mV | -50.80 | [27, 30] |
| $h_2$ | Voltage sensitivity of activation | mV | 6.60 | [27, 30] |
| $h_3$ | $\tau_{\text{hc}}$ basal value | s | $9.18 \times 10^{-2}$ | F [27], Fig. S2 |
| $h_4$ | $\tau_{\text{hc}}$ amplitude | s | 1.10 | F [27], Fig. S2 |
| $h_5$ | Voltage of maximum $\tau_{\text{hc}}$ | mV | -42 | F [27], Fig. S2 |
| $h_6$ | Characteristic width of $\tau_{\text{hc}}$ | mV | 18.50 | F [27], Fig. S2 |
| $g_{\text{hc}}$ | Conductance density | pS $\mu\text{m}^{-2}$ | 193.50 | A<sup>b</sup> [8] |
| $I_{\text{hc}}$ | Current density | fA $\mu\text{m}^{-2}$ | Function | eq. 19 |
| Ca<sub>v</sub> + BK module |  |  |  |  |
| Equilibrium parameters for $V$ and $C$ | | | | |
| $g_L$ | Leakage conductance density | pS $\mu\text{m}^{-2}$ | 1.94 | F [8]<sup>c</sup> |
| $E_L$ | Reversal potential of leakage current | mV | 80.05 | D (eq.3) |
| $\sigma_C$ | Ca<sub>v</sub>-independent Ca<sup>2+</sup> source rate constant | nM s<sup>-1</sup> | $1.14 \times 10^8$ | D (sec. 2.4) |
| $\delta_C$ | Ca<sup>2+</sup> extrusion rate constant | s<sup>-1</sup> | $1.16 \times 10^6$ | MF |
| Ca<sub>v</sub> channels |  |  |  |  |
| $f_{\text{cv}}^o$ | Fraction of open channels | | Variable | eq. 24 |
| $f_{\text{cv}}^c$ | Fraction of closed channels | | Variable | eq. 24 |
| $\alpha_{\text{cv}}$ | Inactivation recovery rate | s<sup>-1</sup> | Function | eq. 23 |
| $\beta_{\text{cv}}$ | Opening rate | s<sup>-1</sup> | Function | eq. 23 |
| $\gamma_{\text{cv}}$ | Inactivation rate | s<sup>-1</sup> | Function | eq. 23 |
| $v_1$ | Inactivation recovery rate constant | mV<sup>-1</sup> s<sup>-1</sup> | 10.10 | MF |
| $v_2$ | inactivation recovery rate voltage inflexion point | mV | -55 | MF |
| $v_3$ | Inactivation recovery rate voltage sensitivity | mV | 1.42 | MF |
| $v_4$ | Opening rate constant | mV<sup>-1</sup> s<sup>-1</sup> | 33.80 | MF |
| $v_5$ | Opening rate voltage inflexion point | mV | -39 | MF |
| $v_6$ | Opening rate voltage sensitivity | mV | 2.10 | MF |
| $v_7$ | Inactivation rate constant | mV<sup>-1</sup> s<sup>-1</sup> | 8.10 | MF |
| $v_8$ | Inactivation rate voltage inflexion point | mV | -18 | MF |
| $v_9$ | Inactivation rate voltage sensitivity | mV | 7.50 | MF |
| $g_{\text{cv}}$ | Conductance density | pS $\mu\text{m}^{-2}$ | 185.16 | MF<sup>d</sup> |
| $I_{\text{cv}}$ | Current density | fA $\mu\text{m}^{-2}$ | Function | eq. 26 |
| BK channels |  |  |  |  |
| $f_{\text{bk}}^o$ | Fraction of open channels | | Variable | eq. 35 |
| $\alpha_{\text{bk}}$ | Opening rate | s<sup>-1</sup> | Function | eq. 36 |
| $\beta_{\text{bk}}$ | Closing rate constant | s<sup>-1</sup> | 97 | MF |
| $b_1$ | Opening rate constant | nM<sup>-1</sup> s<sup>-1</sup> | 0.97 | MF |
| $b_2$ | Opening rate [Ca<sup>2+</sup>]<sub>i</sub> inflexion point $\alpha_{\text{bk}}$ | nM | 316 | MF |
| $b_3$ | Opening rate [Ca<sup>2+</sup>]<sub>i</sub> sensitivity | nM | 30 | MF |

Table A1 – continued from previous page
| Symbol | Description | Units | Reference value | Reference |
| --- | --- | --- | --- | --- |
| $g_{bk}$ | Conductance density | $\text{pS } \mu\text{m}^{-2}$ | 214.50 | MF<sup>e</sup> |
| $I_{bk}$ | Current density | $\text{fA } \mu\text{m}^{-2}$ | Function | eq. 37 |
| **CatSper + sNHE module** |  |  |  |  |
| Equilibrium parameters for $V$ and $C$ | | | | |
| $g_L$ | Leakage conductance density | $\text{pS } \mu\text{m}^{-2}$ | 9 | MF<sup>f</sup> |
| $E_L$ | Reversal potential of leakage current | mV | -20.40 | D (eq. 3) |
| $\sigma_C$ | CatSper-independent $\text{Ca}^{2+}$ source rate constant | $\text{nM s}^{-1}$ | $3.72 \times 10^7$ | D (sec. 2.4) |
| $\delta_C$ | $\text{Ca}^{2+}$ extrusion rate constant | $\text{s}^{-1}$ | $4.27 \times 10^5$ | MF |
| sNHE exchangers |  |  |  |  |
| $f_{nh}^a$ | Fraction of active exchangers | | Variable | eq. 20 |
| $\alpha_{nh}$ | Activation rate | $\text{s}^{-1}$ | Function | eq. 21 |
| $\beta_{nh}$ | Inactivation rate | $\text{s}^{-1}$ | Function | eq. 21 |
| $n_1$ | Activation rate constant | $\text{mV}^{-1} \text{s}^{-1}$ | 1.51 | MF |
| $n_2$ | activation rate voltage inflexion point | mV | -65 | MF |
| $n_3$ | activation rate voltage sensitivity | mV | 7 | MF |
| $n_4$ | inactivation rate constant | $\text{mV}^{-1} \text{s}^{-1}$ | 0.90 | MF |
| $n_5$ | inactivation rate voltage inflexion point | mV | -30 | MF |
| $n_6$ | inactivation rate voltage sensitivity | mV | 7 | MF |
| $J_{\max}$ | Maximal $\text{H}^+$ flux by total active exchangers | $\text{fmol s}^{-1}$ | $3 \times 10^{-7}$ | MF |
| $K_{\text{Na}}$ | $\text{Na}^+$ dissociation constant | mM | 25 | MF [72]<sup>g</sup> |
| CatSper channels |  |  |  |  |
| $f_{cs}^o$ | Fraction of open channels | | Function | eq. 27 |
| $m_{cs}$ | Fraction of channels with voltage- and pH-dependent gate open | | Variable | eq. 28 |
| $h_{cs}$ | Fraction of channels with $\text{Ca}^{2+}$-dependent gate open not inactivated | | Variable | eq. 28 |
| $v_{cs}$ | pH-dependent voltage leading to half-maximal gate opening at equilibrium | mV | Function | eq. 31 |
| $s_1$ | Voltage-dependent gate opening rate constant | $\text{s}^{-1}$ | 270 | MF |
| $s_2$ | $m_{cs}$ voltage sensitivity | mV | 10 | MF [14, 79], Fig. S3 |
| $s_3$ | Amplitude of $v_{cs}$ response to $\text{pH}_i$ | mV | 79.70 | F [75], Fig. S3 |
| $s_4$ | $\text{pH}_i$ leading to half-maximal $v_{cs}$ value | | 7.20 | MF [75], Fig. S3 |
| $s_5$ | Hill-coefficient of $v_{cs}$ | | 35.52 | F [75] |
| $s_6$ | $\text{pH}_i$-independent, minimal $v_{cs}$ value | mV | -41 | MF, Fig. S3 |
| $s_7$ | Scaling factor | nM | 1 | MF |
| $s_8$ | $[\text{Ca}^{2+}]_i$ leading to half-maximal $h_{cs}$ inactivation (IC50) | nM | 700 | MF |
| $s_9$ | Hill coefficient of $\text{Ca}^{2+}$-inactivation curve | | 2.50 | MF |
| $s_{10}$ | $\text{Ca}^{2+}$-inactivation rate constant | $\text{nM}^{-1} \text{s}^{-1}$ | $1.40 \times 10^{-4}$ | MF |
| $g_{cs}$ | $\text{Ca}^{2+}$ conductance density | $\text{pS } \mu\text{m}^{-2}$ | 39.90 | MF<sup>h</sup> [75] |
| $I_{cs}$ | Current density | $\text{fA } \mu\text{m}^{-2}$ | Function | eq. 34 |
| **Common variables and parameters** |  |  |  |  |
| $V$ | Membrane potential | mV | Variable | eq. 1 |
| $E_m$ | Resting membrane potential | mV | -40 | [8, 35, 80, 81] |

Table A1 – continued from previous page
| Symbol | Description | Units | Reference value | Reference |
| --- | --- | --- | --- | --- |
| $C_m$ | Specific capacitance of flagellum membrane | pF $\mu\text{m}^{-2}$ | $1 \times 10^{-2}$ | [82]<sup>i</sup> |
| $v_f$ | Intraflagellar volume | fL | 1.60 | [21] |
| $s_f$ | Surface area of flagellar membrane | $\mu\text{m}^2$ | 30 | D<sup>j</sup> |
| $[\text{K}^+]_o$ | Extracellular potassium concentration | mM | 9 | [83]<sup>k</sup> |
| $[\text{K}^+]_i$ | Resting intraflagellar potassium concentration | mM | 219 | [69] |
| $[\text{Na}^+]_o$ | Extracellular sodium concentration | mM | 423 | [83]<sup>k</sup> |
| $[\text{Na}^+]_i$ | Resting intraflagellar sodium concentration | mM | 20 | [36] |
| $\text{pH}_o$ | Extracellular pH | | 8 | [83]<sup>k</sup> |
| $\text{pH}_r$ | Resting intraflagellar pH | | 7.10 | [36] |
| $[\text{Ca}^{2+}]_o$ | Extracellular calcium concentration | mM | 10 | [83]<sup>k</sup> |
| $C_r$ | Resting intraflagellar calcium concentration | nM | 100 | [10, 36] |
| $C$ | Intraflagellar calcium concentration | nM | Variable | eq. 2.1.4 |
| $H$ | Intraflagellar proton concentration | M | Variable | eq. 5 |
| $\sigma_H$ | sNHE-independent proton source rate constant | $\text{M s}^{-1}$ | $1.50 \times 10^{-7}$ | D (sec. 2.4) |
| $\delta_H$ | Proton extrusion/consumption rate constant | $\text{s}^{-1}$ | 1.28 | D (eq. 5) |
| $E_K$ | $\text{K}^+$ equilibrium potential | mV | −80 | D<sup>l</sup> |
| $E_{\text{hc}}$ | Reversal potential of spHCN current | mV | −18 | [27–29] |
| $E_{\text{Ca}}$ | $\text{Ca}^{2+}$ equilibrium potential | mV | 144 | D<sup>m</sup> |

<sup>i</sup>Conventional value for eukaryotic cytoplasmic membranes<sup>j</sup>Calculated as the lateral surface of a cylinder with length 40 $\mu\text{m}$ and diameter 0.24 $\mu\text{m}$ <sup>k</sup>Typical concentration in artificial seawater used for embryology experiments on marine organisms<sup>l</sup>Calculated with the Nernst equation, $E = \frac{RT}{zF} \ln \frac{[X]_o}{[X]_i}$ , having temperature $T=17^\circ\text{C}$ <sup>m</sup>Calculated with the Nernst equation, setting temperature $T=17^\circ\text{C}$ , and using the $C_r$ parameter as internal concentration value

## Supporting Information

**Fig. S1** Kinetics of intracellular cGMP concentration elicited by different SAP concentrations.

**Fig. S2** Calibration of spHCN gating parameters.

**Fig. S3** Calibration of the CatSper’s pH- and voltage-dependent gate.

